# SUN-MKL1 crosstalk regulates nuclear deformation and fast motility of breast carcinoma cells in fibrillar ECM micro-environment

**DOI:** 10.1101/2020.11.24.396390

**Authors:** Ved P Sharma, James Williams, Edison Leung, Joe Sanders, Robert Eddy, James Castracane, Maja H Oktay, David Entenberg, John S Condeelis

## Abstract

Aligned collagen fibers provide topography for the rapid migration of single tumor cells (streaming migration) to invade the surrounding stroma, move within tumor nests towards blood vessels to intravasate and form distant metastases. Mechanisms of tumor cell motility have been studied extensively in the 2D context, but the mechanistic understanding of rapid single tumor cell motility in the *in vivo* context is still lacking. Here, we show that streaming tumor cells *in vivo* use collagen fibers with diameters below 3 μm. Employing 1D migration assays with matching *in vivo* fiber dimensions, we found a dependence of tumor cell motility on 1D substrate width, with cells moving the fastest and the most persistently on the narrowest 1D fibers (700 nm – 2.5 μm). Interestingly, we also observed nuclear deformation in the absence of restricting extracellular matrix pores during high speed carcinoma cell migration in 1D, similar to the nuclear deformation observed in tumor cells *in vivo*. Further, we found that actomyosin machinery is aligned along the 1D axis and actomyosin contractility synchronously regulates cell motility and nuclear deformation. To further investigate the link between cell speed and nuclear deformation, we focused on the Linker of Nucleoskeleton and Cytoskeleton (LINC) complex proteins and SRF-MKL1 signaling, key regulators of mechanotransduction, actomyosin contractility and actin-based cell motility. Analysis of The Cancer Genome Atlas dataset showed a dramatic decrease in the LINC complex proteins SUN1 and SUN2 in primary tumor compared to the normal tissue. Disruption of LINC complex by SUN1+2 KD led to multi-lobular elongated nuclei, increased tumor cell motility and concomitant increase in F-actin, without affecting Lamin proteins. Mechanistically, we found that MKL1, an effector of changes in cellular G-actin to F-actin ratio, is required for increased 1D motility seen in SUN1+2 KD cells. Thus, we demonstrate a previously unrecognized crosstalk between SUN proteins and MKL1 transcription factor in modulating nuclear shape and carcinoma cell motility in an *in vivo* relevant 1D microenvironment.

## INTRODUCTION

The tumor microenvironment (TME) plays an essential role in breast cancer invasion and metastasis (1–5). TME consists of ECM, stromal cells (e.g. cancer associated fibroblasts, adipocytes) and immune cells (e.g. macrophages, neutrophils etc.) (6–8), and drives breast tumor progression through mechanical and chemical cues (1, 2, 4, 8). The biomechanical properties of tumor ECM (mammographic breast density, ECM stiffness, crosslinking, topography and alignment) play important roles during breast tumor progression. During tumorigenesis, there is an increase in the deposition of matrix proteins, most notably fibrillar collagens, accompanied by extensive matrix remodeling (9–11). Accumulation of collagen I is associated with increased risk of metastasis in breast cancer (12–14).

Striking changes in collagen I architecture (topography and alignment) are observed during breast tumor progression. Multiple studies have reported aligned collagen fibers oriented perpendicular to the tumor boundary in human breast tumor tissue sections (15, 16) and in xenograft breast tumors in mice (17, 18). In contrast, normal breast stroma contains curly collagen fibrils (11, 19, 20). In human samples, collagen fiber alignment correlates with poor disease-specific and disease-free survival (15). Mammographic breast density, and collagen fiber alignment and cross linking are associated with metastasis and poor prognosis in breast cancer (21, 22) and tumor cell dissemination in mouse mammary tumors (23) suggesting an important role for collagen fiber alignment in breast cancer metastasis.

Intravital imaging of mammary tumors has demonstrated that the disseminating tumor cell population moves as single carcinoma cells at high speeds on aligned collagen fibers *in vivo* (5, 24–27). Two-dimensional (2D) assays typically used for tumor cell migration studies lack physiological relevance. For example, 2D assays do not capture elongated cellular/nuclear morphology and high-speed migration behaviors observed *in vivo* (5, 28–33). Our goal in this study was to investigate the single carcinoma cell migration behavior on *in vivo*-like linear collagen fibers. Multiple previous studies have utilized various 1D *in vitro* assays – micropatterned lines, cylindrical fibers, confined microchannels and grooved substrates to model cell migration *in vivo* (28, 29, 34–38). These 1D assays capture many of the cellular characteristics observed *in vivo*, e.g. elongated cellular morphology, rapid cell migration, posterior centrosome position, contractility dependence of migration, and in the case of mammary tumors, alternating linear streams of tumor cells and macrophages (28, 30). Using these 1D and 3D *in vitro* models of cell motility, fiber alignment was found to be a prominent parameter for cell motility (22, 38, 39). However, the relationship of the dimensions of 1D fibers to cell motility phenotype *in vivo* and reconstitution of this relationship *in vitro* has been lacking.

Here, we report the development of an *in vitro* 1D fiber tumor cell motility assay based on the insights obtained from intravital imaging of the modes of cancer cell migration observed in breast tumors and the characterization of mammary tumor ECM fiber dimensions used for this migration *in vivo*. This 1D fiber tumor cell motility *in vitro* assay combined with live imaging of single carcinoma cells led to new insights into the regulators of high-speed tumor cell migration in physiological 1D microenvironments. In particular, we found critical roles of ECM fiber diameter and actomyosin contractility constrained to 1D as dominant factors regulating tumor cell motility and nuclear shape changes. In addition, we found and studied a crosstalk between the LINC complex and SRF-MKL1 signaling on the mechanobiology of tumor cells during high speed cell motility in 1D.

## RESULTS

### Breast carcinoma cells show fast motility primarily on small diameter ECM fibers *in vivo*

In order to reconstitute *in vitro* tumor cell motility with high fidelity to the *in vivo* tumor cell motility phenotype, we first determined the *in vivo* tumor ECM architecture. Using MTLn3 and PyMT mammary tumor models in mice, we imaged the fibrillar collagen associated with tumor cell motility *in vivo* using second harmonic generation (SHG) intravital microscopy (Fig. 1A, B, movie 1). We made 3D reconstructions from the z-stack movies of collagen ECM, and as expected, found that the collagen fibers had a round topography (movie 1). We analyzed 3D reconstruction movies for collagen fiber diameters and found that the most common fiber diameter, in both MTLn3 and PyMT models, was in the 2-3 μm range with the majority being < 3 μm in diameter (Fig. 1C). We also found that tumor cells *in vivo* prefer to move on fibers of < 3 μm in diameter (Fig. 1D).

**Figure 1:**
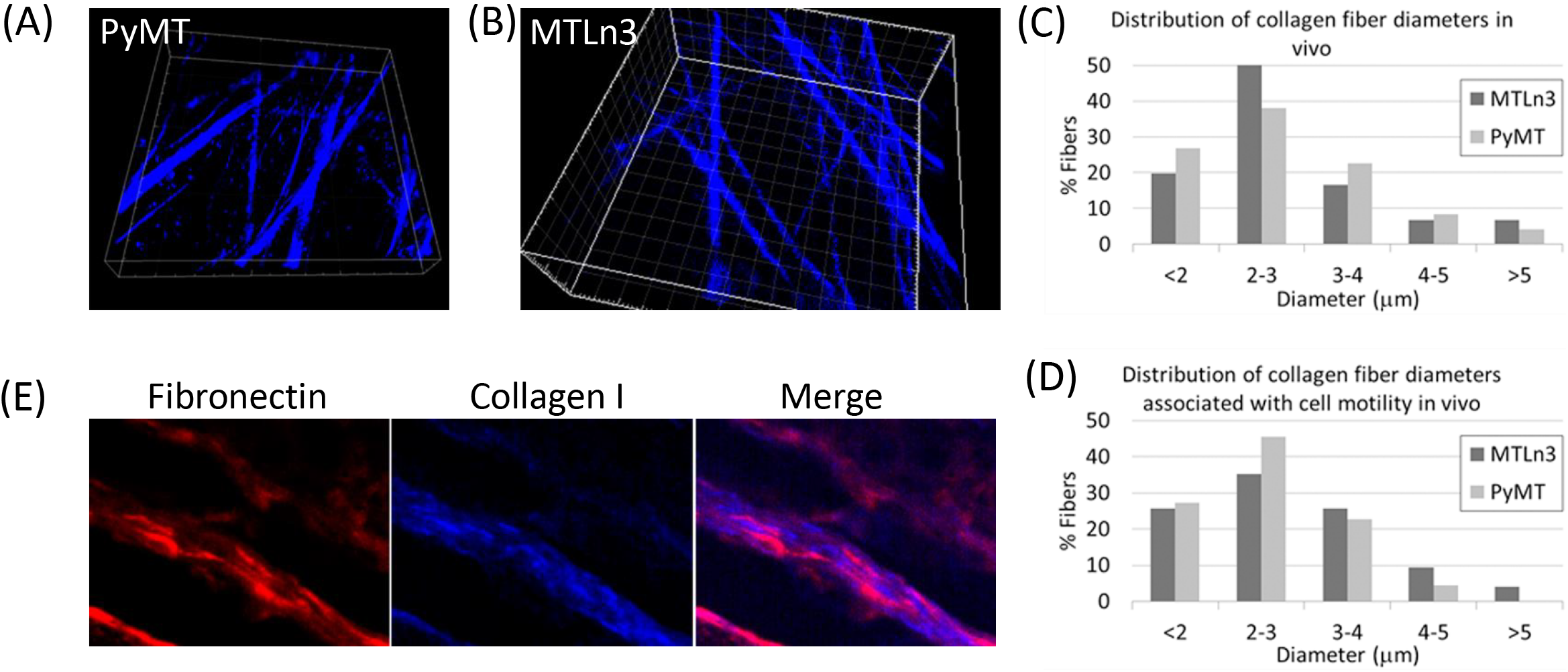
*In vivo* characterization of collagen fiber diameter and composition in mammary tumors. (A, B) Stills from the 3D reconstruction movies of collagen fibers *in vivo* using second harmonic intravital imaging in two mice models - PyMT (A) and MTLn3 (B). (C) Histogram of collagen fiber diameters in PyMT and MTLn3 models. (D) Histogram of tumor cell motility-associated collagen fiber diameters in PyMT and MTLn3 models. (E) Immunofluorescence staining for fibronectin (red, left panel) with second harmonic generation for collagen fibers (blue, middle panel) in MTLn3 tumor sections demonstrates that ECM fibers *in vivo* are composed of both fibronectin and collagen I.

Next, we determined the ECM composition of these *in vivo* fibers. We stained frozen tumor sections with fibronectin antibody and found that ECM fibers are composed of both collagen I (SHG signal) and fibronectin (fluorescence antibody signal). We also checked for other ECM molecules, such as laminin and collagen IV, but did not see any staining in tissue sections (data not shown).

### Fabrication and characterization of *in vitro* 1D fibers

Based on our observations of single breast tumor cell migration on collagen fibers *in vivo* (5, 28, 30) and the fiber diameter characterization we performed above, we set out to create artificial linear fibers matching the *in vivo* ECM fiber diameter and composition. We used electro-spinning technology to generate polymeric fibers made from poly lactic-co-glycolic acid (PLGA) (Fig. 2A). Fibers were electro-spun and suspended over a glass coverslip by attaching them to the PDMS glue at two ends (Fig. 2B). To avoid cell attachment in the 2D area, the glass coverslip was coated with PVA (34) to prevent spreading of tumor cells on the PVA surface (Fig. S1). We performed bright-field microscopy to visualize the fibers and observed single linear fibers (Fig 2C). To visualize fibers at higher resolution, we imaged them using scanning electron microscopy and quantified the fiber diameters (Fig. 2D, E). We found that we can generate fibers with diameters in the nanometer to micrometer range by changing PLGA concentration, with 21% PLGA generating the 2 μm fibers.

**Figure 2:**
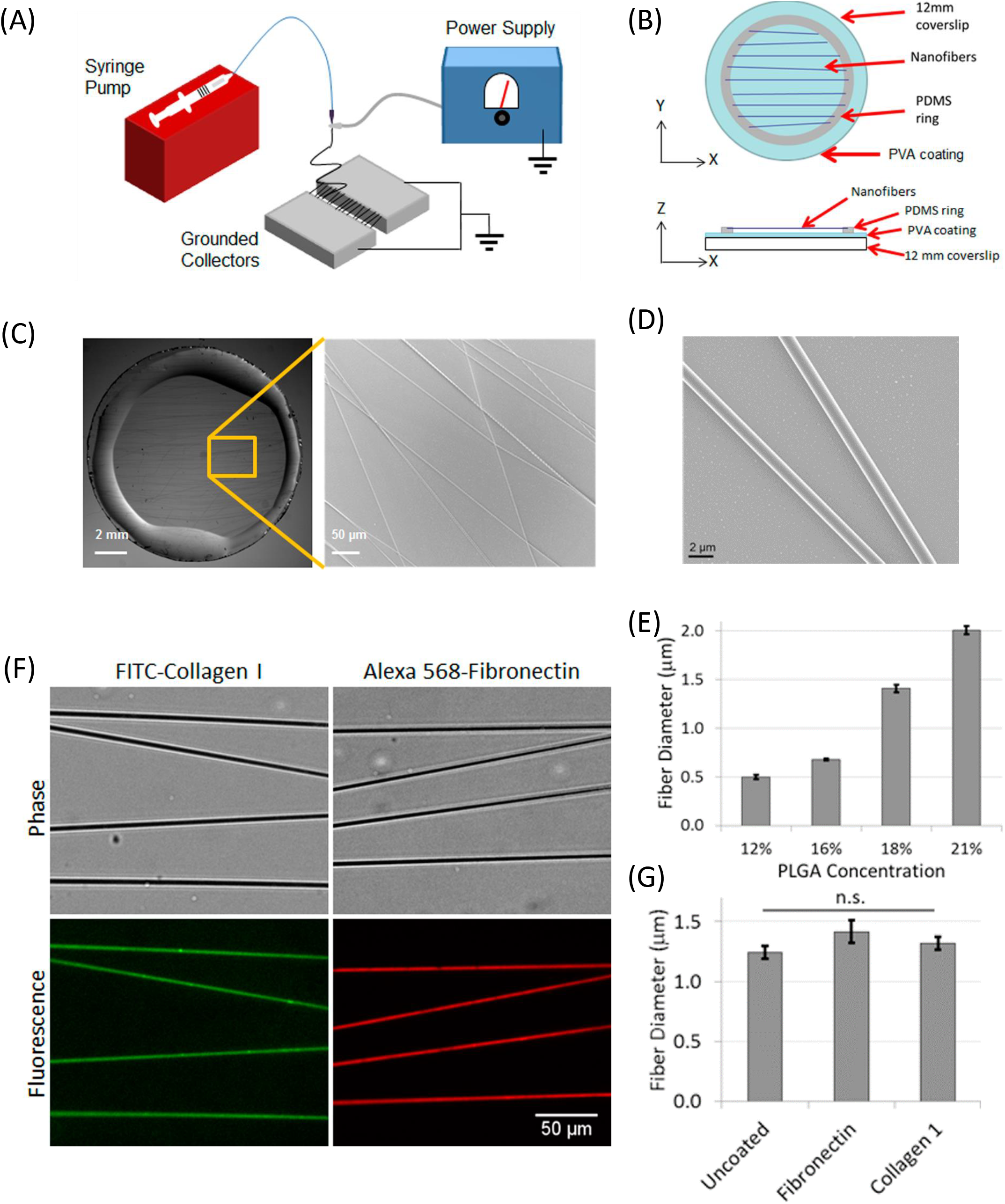
The *in vitro* 1D fiber assay setup: fabrication of PLGA fibers, their size characterization and ECM coating. (A) Schematic of electro-spinning technique setup for the 1D fiber generation. (B) A cartoon depicting the components of 1D fiber assay setup in both X-Y and X-Z views. Fibers are glued at two ends to a PDMS ring, which keeps fibers suspended in air above the PVA-coated 2D surface (cyan). (C) Images of 1D fibers deposited on PDMS glue ring, which keeps fibers suspended in air. Individual fibers can be seen in the zoomed view. (D) Scanning electron microscopy high-resolution image of 1D fibers showing cylindrical morphology of individual fibers. (E) Quantification of PLGA concentration vs. 1D fiber diameter using scanning electron microscopy images of fibers spun at different PLGA concentrations. (F) Images of 1D fibers coated with different ECM molecules (collagen I or fibronectin), showing a uniform and selective coating of 1D fibers with no ECM deposition underneath the fibers on the 2D surface. (G) Quantification of 1D fiber diameter with and without different ECM coatings.

Next, ECM (collagen I or fibronectin) coating conditions were optimized to obtain a uniform ECM coating along the fibers (Fig. 2F). To check the effect of ECM coating on fiber thickness, we performed atomic force microscopy (AFM) on uncoated and ECM coated fibers and found no significant difference in fiber diameter after ECM coating (Fig. 2G).

### Dimensionally-matched physiological 1D substrates mimic *in vivo* fast tumor cell motility

To evaluate the *in vivo* relevance of our 1D fiber assay, we plated mammary carcinoma cells (MTLn3) on ECM coated fibers and performed live imaging (Movie 2). Cells displayed elongated morphology while moving on fibers, similar to what is seen *in vivo*. Another characteristic feature of carcinoma cell migration *in vivo* is the formation of the multicellular streaming pattern containing alternating tumor cells and macrophages (28, 30). Tumor cells and macrophages, when plated together on the ECM-coated fibers *in vitro*, arranged themselves in the alternating streaming pattern seen *in vivo* (Fig. 3A, B, Movie 3). Encouraged by the *in vivo* relevance of our 1D fiber assay, next we carried out a comprehensive investigation of single tumor cell motility characteristics (speed and persistence) on fibers of different diameters and ECM coatings, and compared it with motility parameters of the same cells on a 2D surface of the same ECM (Fig. S3, movie 4). We found that, on 1D fibers tumor cells move twice as fast and with 10 times higher persistence than in 2D (speed: 1.2 vs 0.6 μm/min, and persistence: 0.75 vs 0.07 μm/min-deg in 1D vs 2D, respectively). Interestingly, the choice of ECM (Collagen I or Fibronectin or a combination, collagen I + Fibronectin) did not affect tumor cell motility parameters in 1D (Fig. 3C, D). Since the choice of ECM did not affect tumor cell motility parameters, we chose fibronectin coating for all the subsequent carcinoma cell motility studies in 1D and 2D.

**Figure 3:**
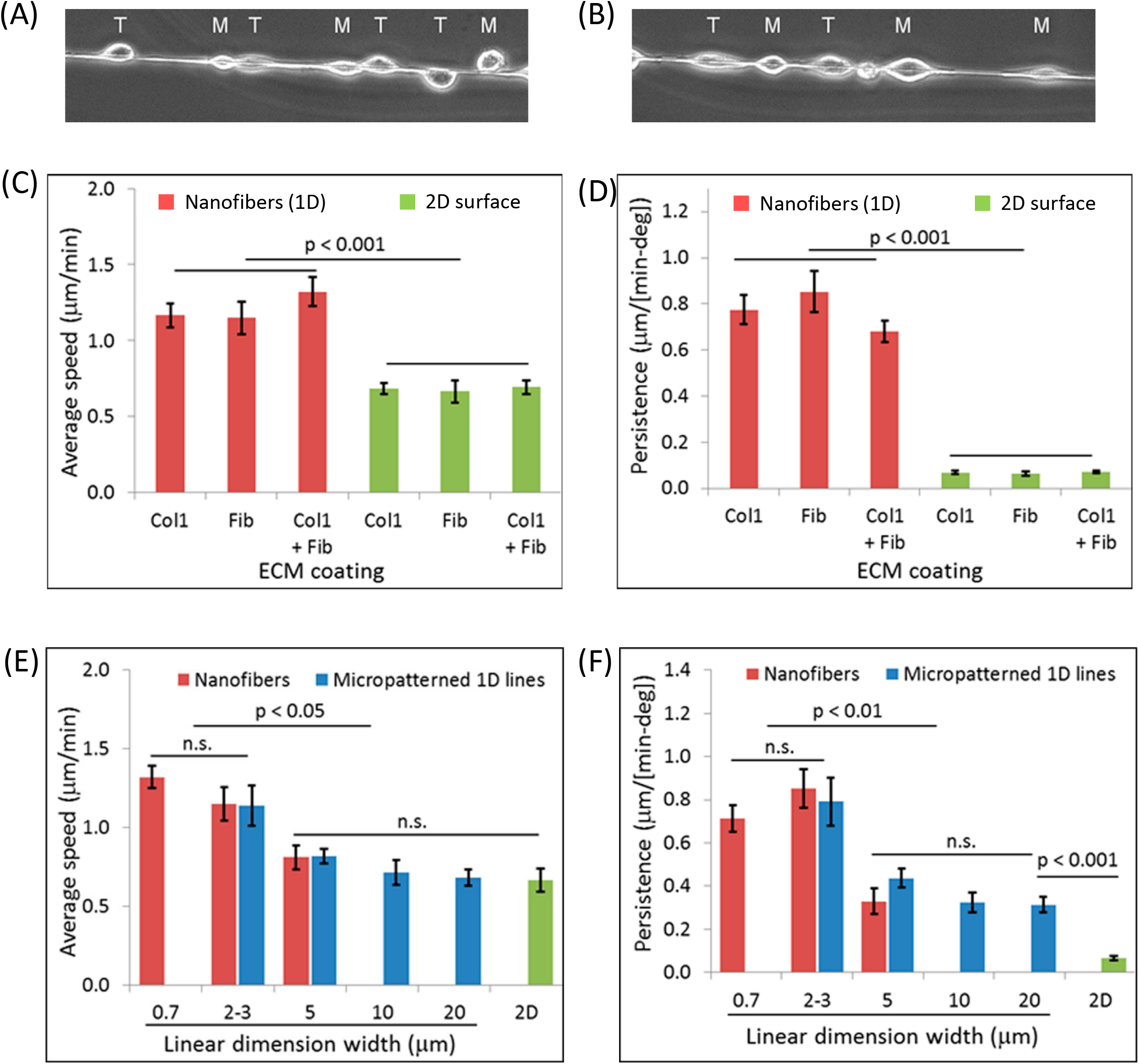
Tumor cells exhibit high-speed migration on thin ECM-coated 1D fibers. (A, B) Assembly of streams composed of alternating cell types is a cell autonomous property on 1D fibers. Two examples of tumor cells (MTLn3, labeled T) and macrophages (BAC1.2F5, labeled M) showing alternating streaming pattern on 1D fibers. (C, D) Quantification of average cell speed (C) and persistence (D) of tumor cells migrating on 2 μm fibers vs on 2D surface coated with different ECM molecules (Collagen I, Fibronectin or Collagen + Fibronectin). Red and green bars represent measurements made on ECM coated (Collagen I, Fibronectin or Collagen I + Fibronectin) 2 μm fibers and on 2D surface, respectively. (E, F) Quantifications of average cell speed (E) and persistence (F) of tumor cells migrating on different thickness 1D fibers (0.7, 2 and 5 μm), micro-patterned 1D lines (2.5, 5, 10 and 20 μm) or 2D surface. Red, blue and green bars represent measurements made on fibronectin coated 1D fibers, 1D micro-patterned lines and 2D surface, respectively.

Next, we examined the effect of varying fiber diameter on tumor cell motility. On smaller diameter fibers (700 nm), tumor cells moved at the same speed and persistence as on 2 μm fibers (Fig. 3E, F). On thicker fibers (5 μm), there was an approximately 30% decrease in cell speed and 60% decrease in persistence, indicating that thicker fibers are not optimum for efficient tumor cell migration. Due to high polymer viscosity at higher PLGA concentrations, we were unable to electrospin fibers > 5 μm in diameter. Therefore, we resorted to using fibronectin-coated micro-patterned 1D lines to study tumor cell motility on > 5 μm 1D lines. First, we compared tumor cell motility parameters on 2.5 and 5 μm thick 1D lines versus 2 and 5 μm thick fibers. We found that tumor cells move at the same speed and persistence on 2.5 and 5 μm 1D lines as they do on 2 and 5 μm fibers, respectively (Fig 3E, F), indicating that micro-patterned 1D lines and 1D fibers can be used interchangeably for tumor cell motility studies. As we increased the micro-patterned line width to 10 and 20 μm, cell speed plateaued out and approached the 2D tumor cell speed (~ 0.6 μm/min). Tumor cell persistence also showed a decreasing trend with increasing 1D line width, although it never reached the 2D persistence value even at 20 μm line width, possibly because the boundary of 1D line restricts cell movement perpendicular to the line axis and contributes toward higher cell persistence, whereas in 2D, cells are free to move in any direction.

### Actomyosin contractility regulates F-actin alignment along 1D axis and high-speed tumor cell motility in 1D

Since actin is the dominant cytoskeleton component contributing toward cell migration (24, 40), we investigated F-actin distribution in tumor cells plated on different width micro-patterned 1D lines and fibers. F-actin stress fibers were primarily oriented along the 1D axis in cells moving on 2 μm fibers or 2.5 μm lines (Fig 4A, B). With increasing 1D line width (5, 10 and 20 μm), stress fibers gradually lost their alignment along the 1D axis and became oriented in all possible directions, approaching the 2D stress fiber alignment distribution (Fig. 4A, B, C). Since higher tumor cell motility parameters correlate with higher F-actin stress fiber alignment along 1D axis, we hypothesize that the aligned F-actin on 2-3 μm 1D lines and fibers helps focus protrusive forces along the 1D axis, leading to high carcinoma cell speed and persistence. In support of this hypothesis, we also found the alignment of focal adhesions along 1D axis in carcinoma cells moving on 2 μm fibers, compared to numerous and often smaller focal adhesions typically seen in cells moving on 2D (Fig. 4D, movie 5).

**Figure 4:**
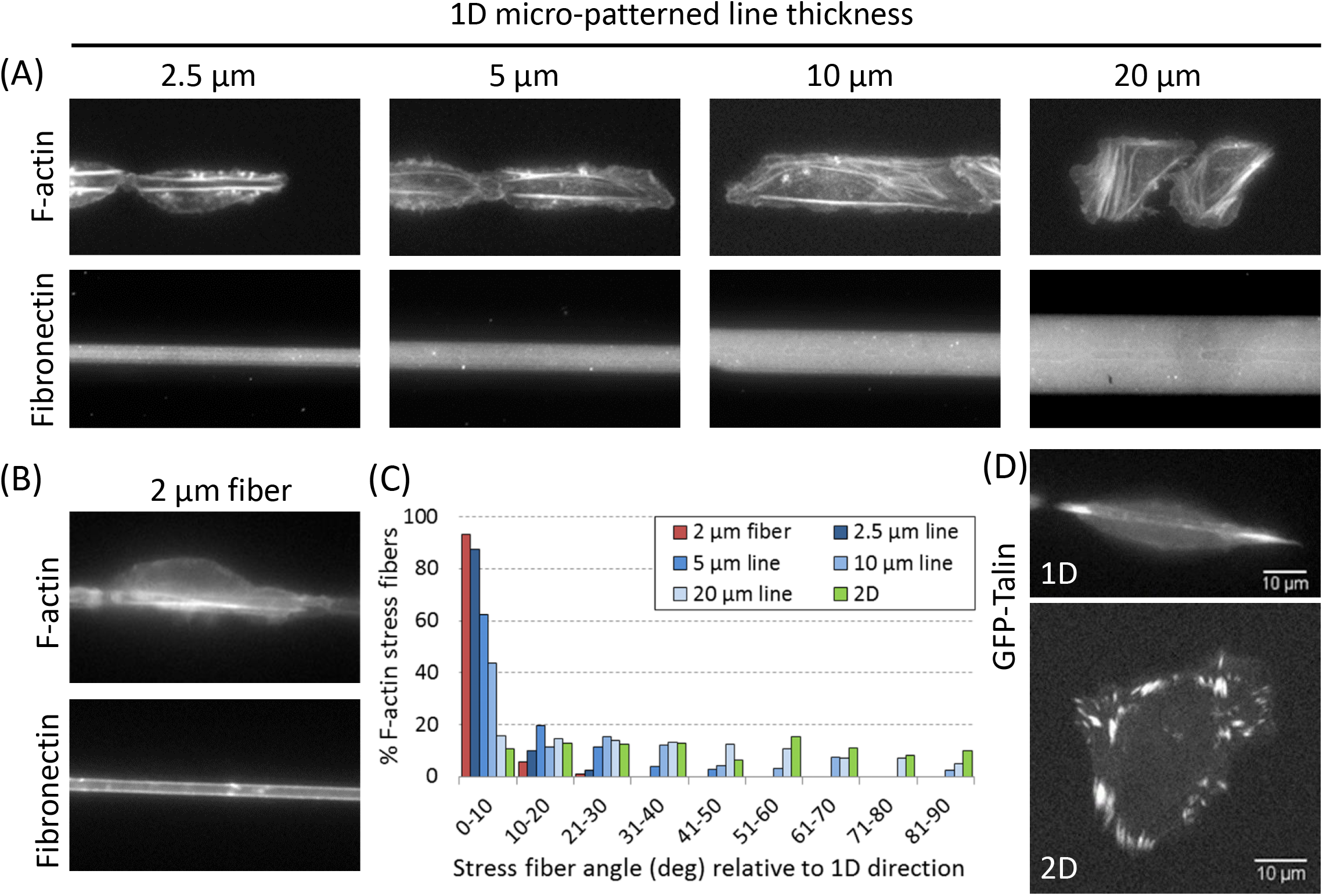
Enhanced alignment of F-actin stress fibers and focal adhesions in tumor cells moving in 1D. (A) F-actin staining in tumor cells moving on different thickness micro-patterned 1D lines (2.5, 5, 10 and 20 μm). (B) F-actin staining in tumor cell moving on 2 μm fiber. (C) Histogram of F-actin stress fiber alignment along different angles. Angles 0 and 90 degrees represent the direction parallel and perpendicular to the 1D axis, respectively. (D) Focal adhesion (GFP-talin) distribution in tumor cells shows aligned focal adhesions in cells moving in 1D (2 μm fibers) vs randomly oriented focal adhesions in cells moving in 2D.

To test the role of aligned F-actin in generating protrusive forces in 1D, we inhibited actomyosin contractility in cells moving in 1D with 5 μM blebbistatin. Using Lifeact as a marker for F-actin in live carcinoma cells, first, we checked F-actin levels in cells before and after the inhibition of actomyosin contractility, and found that there was a sharp decrease in F-actin stress fiber intensity along 1D axis after actomyosin contractility inhibition (Fig 5A, B). This decrease in F-actin intensity after actomyosin contractility inhibition correlated with dramatic decreases in cell speed and persistence (Fig 5C, D, E).

**Figure 5:**
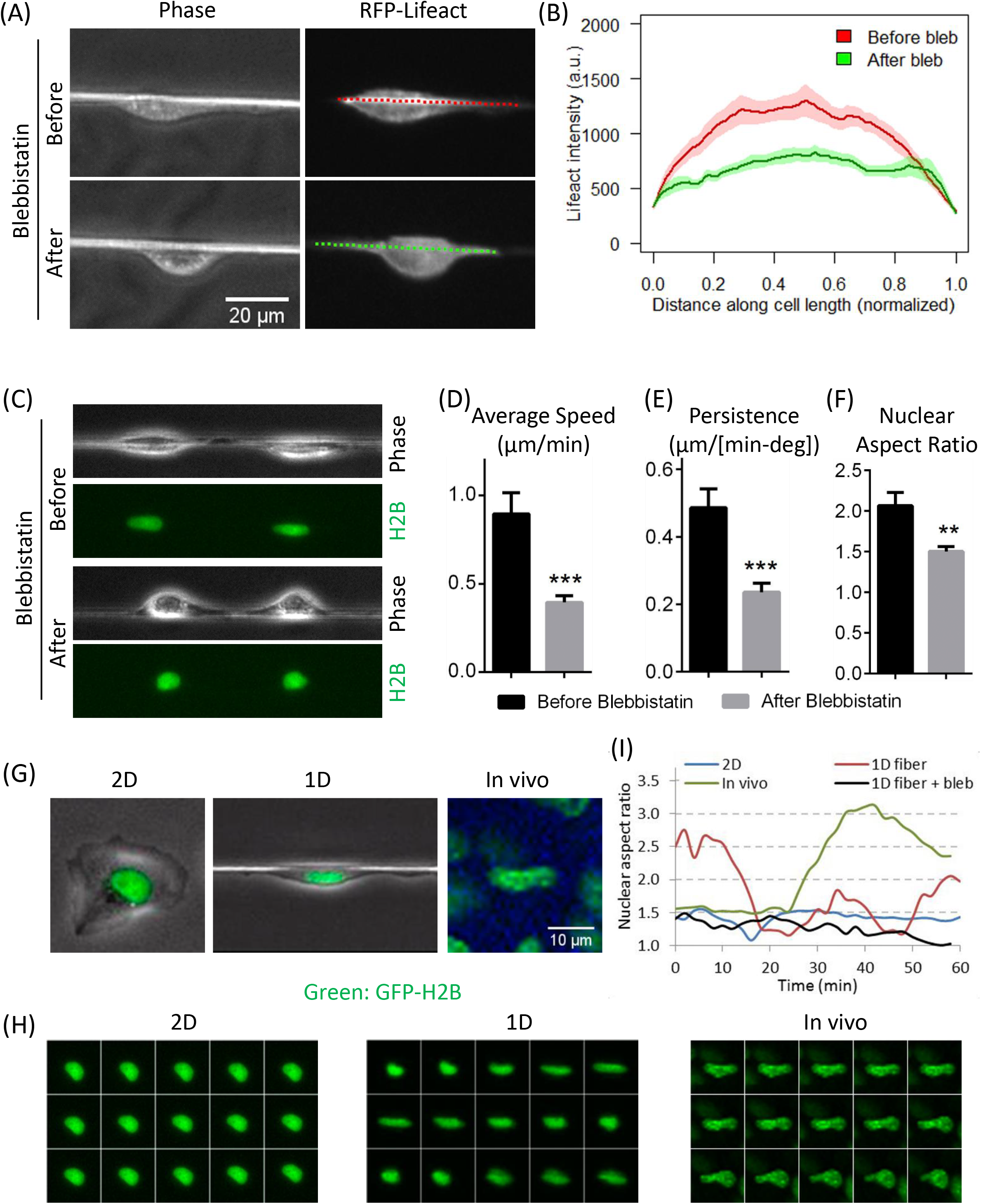
Actomyosin contractility regulates cell-intrinsic nuclear deformation and tumor cell motility in 1D. (A) Phase and RFP channel images of RFP-Lifeact (a marker for F-actin) expressing tumor cells migrating on 2 μm fiber before and after blebbistatin treatment. (B) Quantification of RFP-Lifeact intensity along the cell length (marked with colored dotted lines in figure 6H) migrating on 2 μm fiber before and after blebbistatin treatment. (C) Frames from time-lapse movie 6, showing phase and GFP-H2B channels for MTLn3 cells moving on 2 μm fibers. Mid-way during the movie, cells were treated with 5 μM blebbistatin to inhibit actomyosin contractility. Note the nuclear morphology change from elongated to round after the blebbistatin treatment. (D, E, F) Quantifications of average cell speed (D), persistence (E) and nuclear aspect ratio (F) of tumor cells migrating on 2 μm fibers, before and after blebbistatin treatment (G) Still images from movies 7, 8 and 9 of GFP-H2B MTLn3 cells moving in 2D, 1D (2 μm fibers) and *in vivo*, showing rounder nuclear morphology in 2D vs elongated nuclear morphology in 1D and *in vivo*. (H) Time-lapse images of nucleus (GFP-H2B) shape dynamics, show static rounder nuclear morphology in cells moving in 2D, vs dramatic nuclear shape changes observed in cell moving in 1D and *in vivo*. (I) Plots of nuclear aspect ratio change overtime in cells moving on 2D substrate, *in vivo* and in 1D. Note the actomyosin dependence of nuclear shape change in 1D.

### Fast tumor cell motility and nuclear shape changes are linked together in 1D and *in vivo*, but not in 2D

During our studies on the effect of blebbistatin on cell migration in 1D, we also observed that inhibiting actomyosin contractility led to a rounder nuclear shape compared to the elongated shape observed in untreated cells (Fig 5C, Movie 6). We quantified nuclear aspect ratio and found a significant decrease in nuclear aspect ratio after actomyosin contractility inhibition (Fig 5F). These results suggest that nuclear shape may be intimately linked to the motility potential of tumor cells in 1D, i.e. more elongated the nuclear shape, the higher the motility parameters and vice versa.

To check the link between nuclear shape and cell migration, we investigated the nuclear shape of carcinoma cells in 2D, 1D and *in vivo*. Consistent with high carcinoma cell motility in 1D and *in vivo*, as described above, we found that the nucleus is elongated (i.e. high nuclear aspect ratio) in cells moving in 1D *in vitro* and *in vivo* (Fig 5G). In contrast, the nucleus in cells moving in 2D was found to be round (i.e. low nuclear aspect ratio) (Fig 5G). When we examined nuclear morphology more closely in live movies of GFP-H2B expressing carcinoma cells moving in 1D, we found that nuclear shape showed dynamic elongation and contraction cycles (Fig. 5H, movie 7), compared to static rounder nuclear morphology in cells moving on 2D surface (Fig 5H, Movie 8). Interestingly, we saw similar dynamic nuclear shape changes in the intravital movies of these tumor cells moving *in vivo* (Fig 5H, Movie 9). We quantified the nuclear aspect ratio (major axis/minor axis) and found that in both, *in vivo* and in 1D *in vitro*, the nuclear aspect ratio changes within a similar range (1.5-3.0). In contrast, nuclear aspect ratio in cells moving on 2D surface did not change over time (Fig 5I). These results are surprising, since unlike in an *in vivo* situation where it has been claimed that tumor cells change nuclear shape in order to pass through the narrow ECM pores, cells moving in 1D do not have any pore restriction. These results suggest that tumor cells have the intrinsic ability to modulate nuclear shape, while moving in 1D. Unlike fibroblasts, where a perinuclear actin cap and formin FMN2 have been proposed to regulate cell migration and nuclear shape (41–43), carcinoma cells used in this study do not rely on perinuclear actin cap or FMN2 for cell motility (Figs S4, S5).

### Down-regulation of LINC complex proteins, SUN1 and SUN2, in cancer patients

Irregular nuclear shape and nuclear softness is seen in human laminopathies (a collection of human diseases including certain types of muscular dystrophies, dilated cardiomyopathy, and the premature aging disease Hutchinson–Gilford progeria syndrome) (44). These diseases are primarily caused by the mislocalization or loss of nuclear envelope proteins. The Linker of Nucleoskeleton and Cytoskeleton, or LINC complex, comprising of Nesprin and SUN family proteins of the nuclear envelope, performs a critical role of providing a physical link between the cell cytoskeleton and nucleus. Through this physical connection, LINC complex proteins regulate cellular mechanotransduction. Several studies have reported defects in cell motility after LINC complex disruption (45–47), however the role of LINC complex in carcinoma cell motility in physiological environments is unknown.

The LINC complex, Nesprin and SUN family proteins, reside in the outer and inner nuclear membranes, respectively (48–50). Together they provide a physical link between the cellular cytoskeleton and nuclear lamina and have been implicated in nuclear mechano-transduction and cell motility (45, 50, 51). Global loss of LINC complex components has previously been reported in breast cancer (52). Analysis of The Cancer Genome Atlas (TCGA) dataset further confirmed that the expression of SUN1 and SUN2 is down-regulated in breast cancer compared to normal breast tissue (Figs. 6A, B). Moreover, down-regulation of SUN1 and SUN2 was observed across tumor types (Figs. S6A, B).

**Figure 6:**
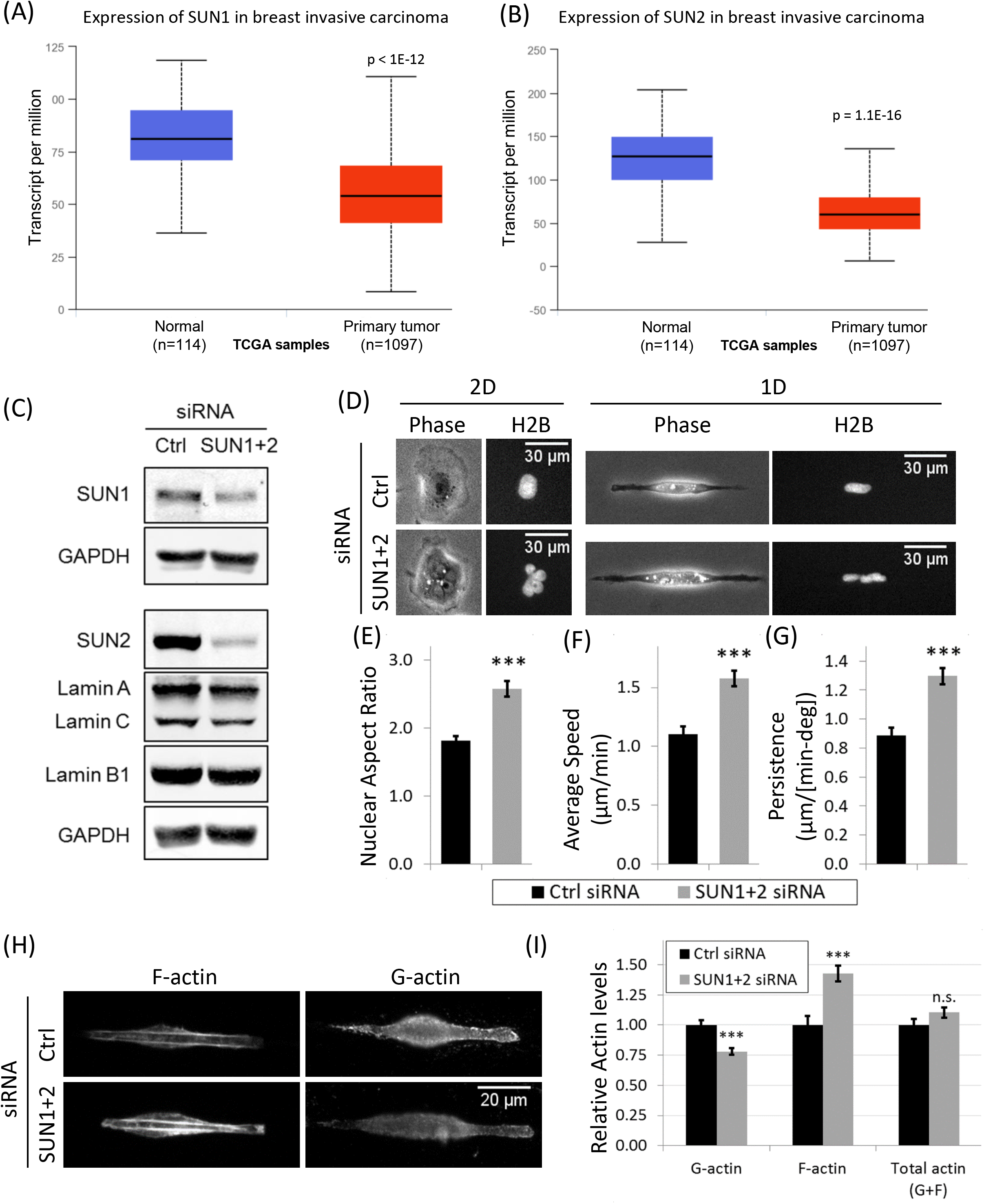
SUN1 and SUN2 down-regulation in breast cancer patients; LINC complex disruption increases actin polymerization and tumor cell motility in 1D, but not in 2D. (A) Box plot analysis shows downregulation of SUN1 in primary breast tumor (n=1097) versus normal breast tissue (n=114) derived from RNA-Seq expression data from TCGA samples. (B) Box plot analysis shows downregulation of SUN2 in primary breast tumor (n=1097) versus normal breast tissue (n=114) derived from RNA-Seq expression data from TCGA samples. (C) Western blots showing SUN1 and SUN2 knockdowns in SUN1 siRNA + SUN2 siRNA treated tumor cells. Blots were also stained with Lamin A/C and Lamin B1 antibodies to check Lamin levels after SUN1+2 KD. (D) Images showing cellular (phase panels) and nuclear (GFP-H2B panels) morphologies after SUN1+2 KD in 1D (2.5 μm micropatterned line) and 2D. Note the multi-lobular nuclear morphology after SUN1+2 KD in both 1D and 2D. (E, F, G) Quantifications of nuclear aspect ratio (D), average tumor cell speed (E) and tumor cell persistence (F) after SUN1+2 KD in 1D. (H) Images showing phalloidin (F-actin) and G-actin staining in control and SUN1+2 KD cells migrating in 1D. Images were acquired at the same exposure time and ND filter settings for each channel. (I) Quantifications of whole cell F-actin, G-actin and total actin (F-actin + G-actin) levels in control and SUN1+2 KD cells migrating in 1D.

### Disruption of LINC complex leads to increased tumor cell migration and actin polymerization in 1D

To further investigate the link between SUN proteins, nuclear shape and motility in carcinoma cells, we disrupted LINC complex by knocking down SUN proteins (SUN1 and SUN2). Using smartpool siRNAs for SUN1 and SUN2, we efficiently knocked down SUN1 and SUN2 (referred to SUN1+2 KD hereafter) in tumor cells (Fig 6C). Single cell immunofluorescence imaging confirmed depletion of SUN1 and SUN2 from the nuclear envelope after SUN1+2 KD (Figs. S6C, D). We investigated nuclear morphology in SUN1+2 KD tumor cells, and found that cells showed irregular nuclear morphology with multiple lobes, folds and membrane invaginations in both 1D and 2D (Fig. 6D, S6C-E, movies 10 and 11), indicative of compromised nuclear mechanotransduction due to defects in nucleo-cytoskeleton integrity. Since Lamins have been shown to regulate nuclear envelope integrity (40, 44), we checked Lamin levels after SUN1+2 KD, and found that Lamin A/C and Lamin B1 levels did not change much after SUN1+2 KD (Fig. 6C). Single cell immunofluorescence imaging also confirmed no change in nuclear envelope localizations of Lamin A/C and Lamin B1 after SUN1+2 KD (Fig S6C-E), i.e. Lamins stay at the nuclear envelope even in the irregularly shaped nucleus after SUN1+2 KD. These results indicate that the nuclear morphology defects observed in SUN1+2 KD cells are not due to off-target effects, such as Lamins. Consistent with the nuclear phenotype, nuclear aspect ratio increased after SUN1+2 KD (Fig 6E).

Next, we investigated the motility parameters of tumor cells in 1D after SUN1+2 KD. Surprisingly, we found that both speed and persistence increased significantly after SUN1+2 KD (Figs. 6F, G). This again suggests that carcinoma cell migration in 1D is linked to nuclear shape, as seen above (Fig 5). Since F-actin is the dominant cytoskeleton component regulating tumor cell migration (24, 40), we investigated the changes in F-actin and G-actin (with specific antibody JLA20, see materials and methods and Fig. S6F) levels in single tumor cells after SUN1+2 KD. We observed a significant increase in F-actin levels and a concomitant decrease in G-actin levels after SUN1+2 KD (Figs. 6H, I) with no change in total actin levels after SUN1+2 KD (Fig. 6I), indicating an upregulation in actin polymerization (G-actin to F-actin conversion). In cells moving on 2D substrates, no changes in tumor cell speed, persistence and actin polymerization (Figs. S6G, H) were observed after SUN1+2 KD, indicating 1D-specific changes in cell motility and associated actin forces. Together, these results demonstrate that LINC complex disruption by SUN1+2 KD leads to 1D-specific increases in actin polymerization and tumor cell motility.

### MKL1 nuclear translocation regulates increased tumor cell 1D motility in SUN1+2 KD cells

We found decreased G-actin levels in SUN1+2 KD cells (Figs. 6H, I). Myocardin-related transcription factor A (MRTF-A), also known as megakaryoblastic leukemia factor 1(MKL1) shuttles to the nucleus from the cytoplasm in response to changes in G and F-actin levels in the cell, where MKL1 binds serum-response factor (SRF) and initiates transcriptional programs for increased actin polymerization and cell motility (53, 54). In particular, transcription factor, MKL1 is known to translocate from the cytoplasm to the nucleus after reduced G-actin levels (55). Therefore, we investigated if MKL1-SRF signaling pathway plays any role in tumor cell motility in 1D. First, we checked if SRF-MKL1 signaling is relevant in our mammary tumor cells by looking at MKL1 distribution in mammary tumor cells before and after serum stimulation (55). We stained serum-starved and serum-stimulated cells with MKL1 antibody. Compared to the cytosolic MKL1 distribution in serum-starved cells, we found a robust nuclear MKL1 localization in serum-stimulated cells (Fig 7A, B), indicating that MKL1-SRF pathway is active in mammary carcinoma MTLn3 cells. Since SUN1+2 KD led to a decrease in G-actin in cells moving in 1D (Figs. 6H, I) and the drop in G-actin levels is known to cause MKL1 translocation from the cytosol to the nucleus (55), we hypothesized that SUN1+2 KD will lead to MKL1 nuclear translocation. We treated tumor cells with control or SUN1+2 siRNA, and plated them on 1D or 2D substrates. MKL1 antibody staining of cells showed nuclear MKL1 translocation after SUN1+2 KD in both 1D and 2D. We quantified nuclear MKL1 signal and found that there was nearly 50% increase in nuclear MKL1 signal after SUN1+2 KD in cells moving in 1D. The increase in nuclear MKL1 signal in cells moving in 2D, though significant, was rather modest (~ 15%). These results indicate that loss of SUN proteins leads to nuclear MKL1 translocation, activating MKL1-SRF gene transcription machinery, leading to increased actin polymerization (Fig. 6H, I) and increased tumor cell 1D motility (Figs. 6 F, G).

**Figure 7:**
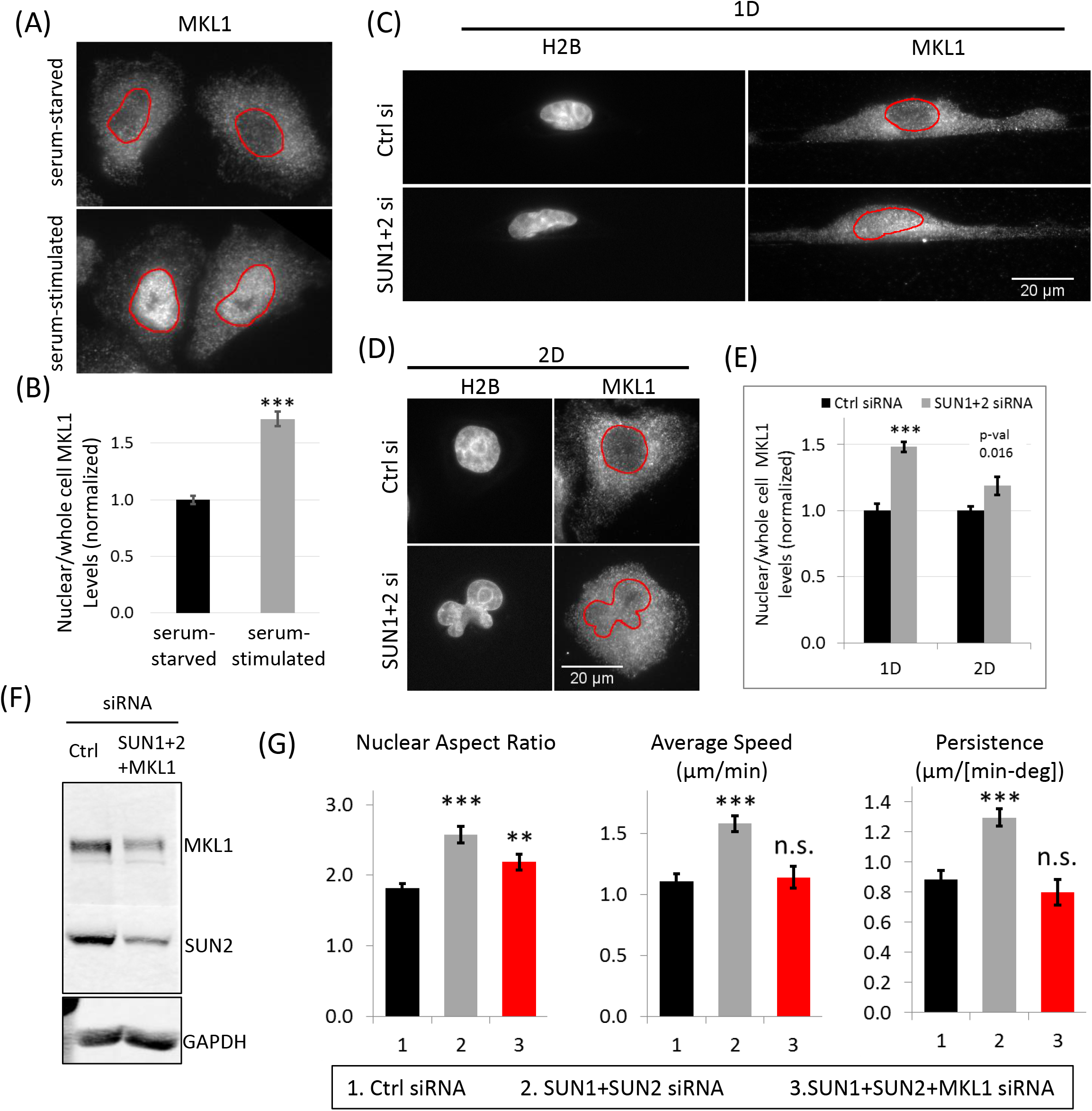
MKL1 is required for increased 1D tumor cell migration after SUN 1+2 KD. (A) MTLn3 carcinoma cells were stained with MKL1 antibody in serum-starved or serum stimulated conditions in 2D. Images show robust MKL1 nuclear translocation after serum stimulation (5% FBS for 5 min). Red outlines show nucleus boundary. (B) Quantification of nuclear/ whole cell MKL1 levels in serum-starved or serum-stimulated MTLn3 carcinoma cells. (C) Images of control siRNA or SUN1+2 siRNA treated H2B-MTLn3 cells migrating on 2.5 μm micropatterned 1D lines, stained with MKL1 antibody to show the nuclear MKL1 translocation after SUN1+2 KD. Red outlines show nucleus boundary. (D) Images of control siRNA or SUN1+2 siRNA treated H2B-MTLn3 cells migrating in 2D, stained with MKL1 antibody to show the nuclear MKL1 translocation after SUN1+2 KD. Red outlines show nucleus boundary. (E) Quantification of nuclear/ whole cell MKL1 levels in control or SUN1+2 siRNA treated cells migrating in 1D or 2D. (F) Western blots showing knockdowns of SUN2 and MKL1 in tumor cells treated with either control siRNA or SUN1+SUN2+MKL1 siRNAs. (G) Quantifications of nuclear aspect ratio, average tumor cell speed and tumor cell persistence after SUN1+SUN2+MKL1 KD in cells migrating in 1D. Control and SUN1+2 siRNA bars are shown here from figure 6E-G for comparison.

Next, we asked if MKL1 signaling is required for increased 1D motility of SUN1+2 KD cells. We treated tumor cell with either control siRNA or siRNAs for SUN1, SUN2 and MKL1. Western blot analysis showed nearly 80% KD of both MKL1 and SUN2 proteins (Fig 7F). We plated triple KD cells (SUN1+2 and MKL1) on 1D or 2D substrates and made time-lapse movies. We quantified nuclear aspect ratio, speed and persistence and found that MKL1 is required for high-speed 1D motility of SUN1+2 KD cells (Fig 7G). MKL1 KD, though, was not able to restore the nuclear morphology to what is seen in control cells (Fig 7G), indicating that MKL1 acts downstream of SUN1+2 mediated nuclear shape change. The effects of MKL1 on tumor cell motility were specific to 1D, as we did not see any change in tumor cell 2D motility after SUN1+2 and MKL1 KDs (Fig S6I).

## DISCUSSION

In this work, we have evaluated the roles of cell-extrinsic (collagen fiber diameter) and cell-intrinsic (SUN and MKL1 signaling) factors influencing tumor cell motility in 1D, the fibrillar ECM microenvironment which tumor cells encounter *in vivo* to invade locally and migrate to the nearest blood vessel for hematogenous dissemination to secondary sites (5, 15, 56). Our work demonstrates that tumor cells move the fastest on 2-3 μm diameter 1D fibers. 1D high-speed migration of tumor cells is closely linked to cell-intrinsic capacity for nuclear deformation. In addition, LINC complex disruption in tumor cells leads to even more enhancement in cell motility in 1D (Fig 8). Interestingly, these results were specific to 1D because tumor cells in 2D not only moved significantly slower, but also showed no changes in 2D motility after SUN and MKL1 signaling perturbations. Highly efficient high-speed tumor cell migration in 1D is achieved through aligning cytoskeletal forces in the tumor cell along the ECM fiber axis, whereas in 2D, cytoskeletal forces are randomly distributed leading to lower net forces in the direction of movement (Fig 8).

**Figure 8:**
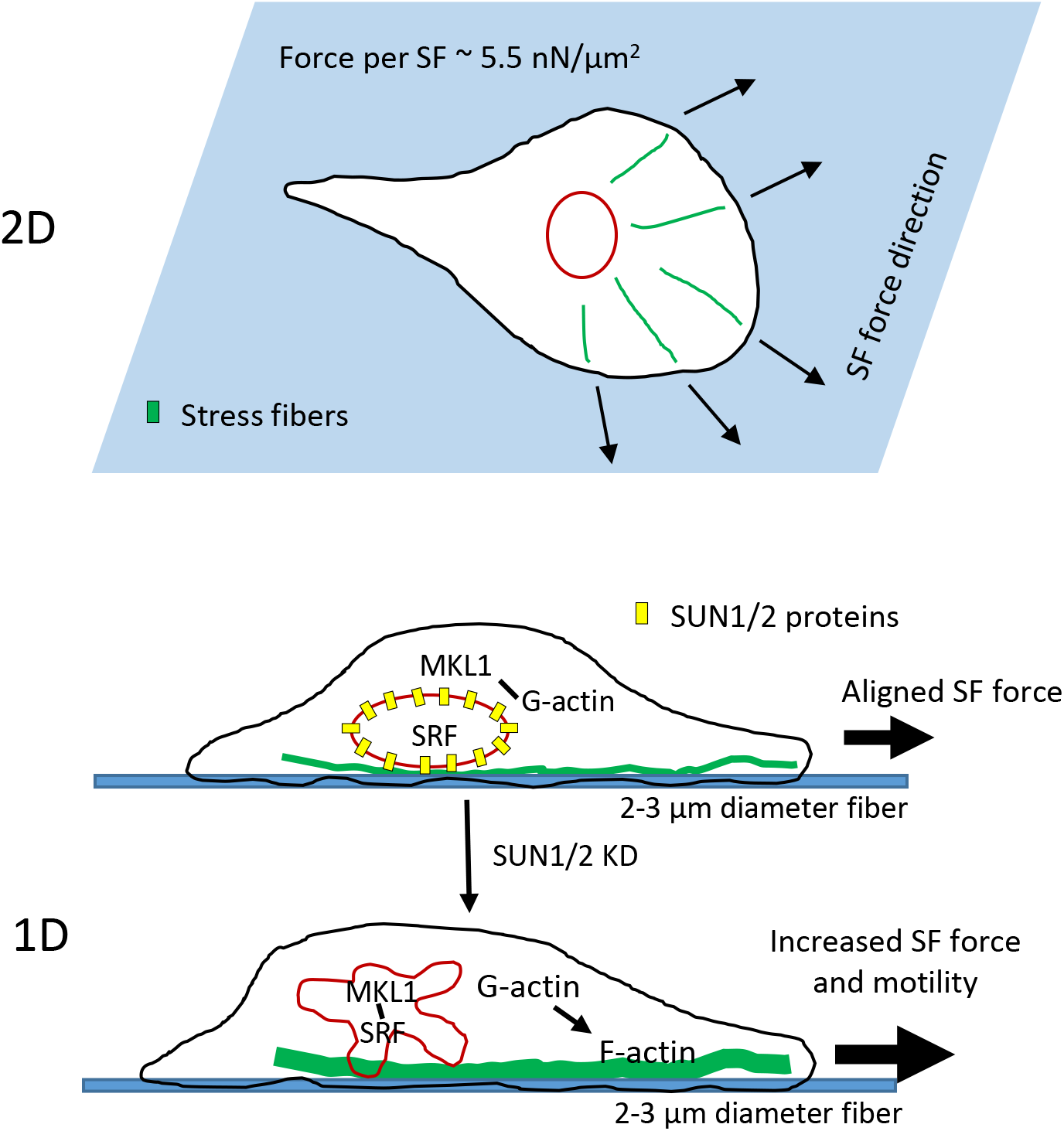
Model of tumor cell migration in fibrillar tumor microenvironment *in vivo*. During invasion, single tumor cells migrate along thin collagen fibers (2-3 μm in diameter) with high speeds. As opposed to 2D environment, where cell spreads leading to distributed actomyosin forces (force per stress fiber ~ 5.5 nN/μm2 (88, 89)), in confined 1D environment, greater aligned actomyosin forces along the fiber 1D axis generate high-speed migration. This 1D high-speed migration is couple to cell-intrinsic nuclear deformation even in the absence of any restricting ECM pores. Downregulation of LINC complex proteins (SUN1 and SUN2) in tumor cells lead to MKL1 nuclear translocation and increased SRF-MKL1 transcriptional activity, which upregulates genes for actin polymerization activity and further enhances tumor cell motility in 1D.

### Collagen fiber diameter is a key regulator of tumor cell motility *in vivo*

The tumor microenvironment ECM presents many biomechanical cues (ECM stiffness, crosslinking, topography and alignment) to sustain high-speed tumor cell motility during tumor progression *in vivo*. In this study, we investigated the role of one such cue, the collagen fiber diameter, on tumor cell motility *in vivo*. Previous studies have imaged ECM collagen fibers *in vivo* in a variety of tissues (rat tail, ovarian tissue, mouse tumor xenografts etc.) and reported their alignments (57–59). A few studies have looked at collagen fiber diameters in healthy tissues and reported to be in the range of 500 nm to a few micrometers (60, 61). To our knowledge, we are the first to report collagen fiber diameter distribution in mammary tumors *in vivo*.

In this study, we have reconstituted with high fidelity the high speed and persistence of streaming migration in tumor cells seen *in vivo* using synthetic fibers and micropatterned tracks composed of physiological ECM. The phenotypes of tumor cells moving on 1D fibers *in vivo*, including the formation of alternating patterns with macrophages and high speed persistent migration with pulsating nuclear deformation were exactly recreated *in vitro* and were fiber diameter dependent. Both micropatterned channels and fibers gave equivalent results and showed the same dependence of cell phenotype on the diameter of the linear ECM, indicating that the diameter and not curvature of the linear ECM dominated cell phenotype.

Previous studies utilizing 1D assays to model motility of different cell types (breast tumor, epithelial, fibroblast, myoblast) used 1D fiber diameters/line widths which were not thin (20-170 μm (62), 20 μm – 2D (36)) or used a narrow range of diameters/widths (0.7-0.8 μm (17), 0.4-1.2 μm (63)). In general, these studies did not find changes in single cell speed as a function of varying 1D width. The Yamada group, on the other hand, used a range of micropatterned line widths (1 - 40 μm) to study fibroblast speed (64). Interestingly, our results on the effect of varying 1D width (0.7 μm – 20 μm) on carcinoma cell speed are in close agreement with their fibroblast speed values, suggesting that extrinsic biomechanical cues from the microenvironment, rather than driver mutations and/or differentiated state of the cell, dominates motility phenotype in both normal and cancer cells.

### Nuclear shape changes are linked to high-speed tumor cell 1D motility and do not require ECM pore restriction

Consistent with previous 1D studies (41, 65, 66), we observed elongated nuclear morphology in tumor cells migrating in 1D (Fig. 5G). Elongated nuclear morphology was further confirmed in carcinoma cells moving *in vivo* (Fig. 5G). One mechanism of nuclear deformation is the compressive lateral actomyosin forces acting on both sides of the nucleus (66). Accordingly, we saw alignments of F-actin and focal adhesions along the 1D axis (Fig. 4). Moreover, inhibition of actomyosin contractility led to rounding of the nucleus in 1D (Fig. 5C, F). Another mechanism for regulating nuclear shape is the apical F-actin caps (41, 42). Unlike fibroblasts, we did not see any apical F-actin caps in carcinoma cells (Fig S4), indicating that lateral actomyosin forces are the primary mechanism behind nuclear elongation in carcinoma cells in 1D.

Previous literature suggests that tumor cells deform nucleus in order to squeeze through small ECM pores encountered during invasion and metastasis (40, 67–70). Here, we show that the tumor cell nucleus is inherently plastic and can display changes in shape in 1D, even in the absence of any restricting ECM pores. The amplitude of nuclear shape change observed in 1D was remarkably similar to nuclear shape changes observed in these cells *in vivo* (Fig. 5H, I). These results suggest that rapid nuclear shape change is linked to high-speed tumor cell migration in 1D, since the inhibition of actomyosin contractility led to the disappearance of nuclear shape change cycles (nucleus became round) and greater than 50% reductions in cell motility parameters. Therefore, nuclear plasticity is not only required for passing through small pore openings in the ECM, but it is also linked to high-speed tumor cell migration in 1D, via the nucleo-cytoskeleton link.

Nuclear elongation is one of the characteristic features of migrating cells *in vivo* (40, 67, 71), which has been confirmed by many 1D studies (38, 41, 42, 65, 66, 72). The nucleus is the largest and stiffest organelle in the cell and it’s been suggested to be an impediment for cell migration *in vivo* (67, 70, 73). In contrast to the nucleus in normal cells, the nucleus in tumor cells is softer (74–78) and sometimes, has aberrant morphology indicated by nuclear folds, invaginations and blebs (79, 80). This allows tumor cells to modulate nuclear shape and escape through narrow openings frequently encountered during tumor cell invasion and metastasis.

### Role of LINC complex and MKL1-driven gene transcription in high-speed tumor cell motility in 1D

The effect of LINC complex disruption on cell motility in 1D-like environments, to our knowledge, has not been previously reported. Using 2D substrates, previous studies (45–47) have reported reductions in cell motility after LINC complex disruption in fibroblast and endothelial cells. However, in the tumor cells studied here, we did not see any changes in 2D cell motility after SUN1+2 knockdown (Fig. S6G). Interestingly, the same tumor cells showed 50% higher motility after SUN1+2 knockdown when moving in 1D environment (Fig. 6F). Previous studies have shown that disrupting nuclear lamins lead to nuclear softening (81, 82). In SUN1+2 knockdown tumor cells, we saw irregular shaped nuclei with nuclear envelope folds, invaginations and blebs (Fig. 6D, movies 10 and 11). These irregular nuclear features are often seen in cancer cells from patients and in normal cells with reduced Lamin protein expression (40, 44). We therefore speculate that SUN1+2 knockdown leads to a softer nucleus in carcinoma cells, allowing them to squeeze through tight spaces encountered during invasion and metastasis. The increased nuclear flexibility of SUN1+2 KD carcinoma cells to pass through narrow ECM constrictions combined with their increased motility in 1D make these cells super invaders during mammary tumor invasion. Two previous studies, one utilizing a 3D transwell migration assay (73) and the other microfluidics (69), which mimic *in vivo* environment more closely than 2D assay, support our findings - that the nucleus is an impediment to cell migration in 3D/1D like environments and that softening the nucleus leads to higher cell motility.

Forces required for increased motility of SUN1+2 knockdown cells are generated by increased F-actin polymerization (Fig. 6H, I), suggesting upregulation of genes involved in F-actin polymerization. Changes in G and F–actin levels in cells are known to cause nuclear translocation of MKL1 (55). The activation of the SRF-MKL pathway upon nuclear translocation targets the transcription of many genes involved in the regulation of actin polymerization, dynamics and protrusion (e.g. cofilin, WASP, Arp2/3), cell-cell and cell-ECM adhesion (e.g. integrin, vinculin and cadherin), enhancing cell motility through a positive feedback regulation of actin polymerization (53, 54, 83). Our novel observation that LINC complex disruption leads to increased MKL1 nuclear translocation suggests cross-talk between LINC complex and SRF-MKL pathway and an important role of SRF-MKL driven gene transcription in 1D tumor cell motility. In fact, using triple knockdowns (SUN1, SUN2 and MKL1) in tumor cells, we show that MKL1 is required for enhanced 1D motility of SUN1+2 KD cells (Figs. 7F-G). Future studies are required to identify specific SRF-MKL transcriptional genes involved in tumor cell motility in physiological environments.

## AUTHOR CONTRIBUTIONS

VPS, JW, RE, JC, MHO, DE, JSC designed research; VPS and EL performed all biological experiments; JW, JS, JC fabricated and characterized 1D fibers; VPS wrote cell tracking ImageJ macro and analyzed all biological data; VPS, MHO, JSC wrote the manuscript; and all authors edited and approved the manuscript.

## ACKNOWLEDGMENTS

We thank Yarong Wang for providing Dendra2-H2B stable MTLn3 cell line and Louis Hodgson for MEF cell line; Frank Macaluso and Leslie Gunther-Cummins (Analytical Imaging Facility, Albert Einstein College of Medicine) for SEM imaging of 1D fibers and Gruss Lipper Biophotonics Center at Albert Einstein College of Medicine for microscopy help, and members of the Condeelis, Segall, Cox, and Hodgson laboratories for guidance. This work was funded by NIH grants CA150344, CA216248, the Gruss Lipper Biophotonics Center and its Integrated Imaging Program and R01 CA 216248-01.

## MATERIALS AND METHODS

### Cell Culture, Antibodies, Immunofluorescence

Rat mammary adenocarcinoma, MTLn3 cells were cultured in α-MEM media, supplemented with 5% FBS and antibiotics as described earlier (28, 29). Mouse embryonic fibroblasts (MEF) were a kind gift from Louis Hodgson. Cells were transfected with Dendra2-H2B and FAC sorted to generate stable cells. Cells were subsequently maintained in G418 to maintain H2B expression. Following antibodies were used: mouse anti-fibronectin antibody (Abcam #6328); mouse anti-G-actin (JLA20) (84) and mouse anti-actin(AC-15), SUN1, SUN2, Lamin A/C, Lamin B1, GAPDH, MKL1 (Cell Signaling). For immunofluorescence experiments cells were fixed in 4% paraformaldehyde, permeabilized in 0.1% Triton-X and blocked in 1% BSA + 1% FBS containing PBS before proceeding with primary and secondary antibodies. For G-actin staining, cells were fixed and permeabilized in cold acetone on ice for 5 min according to (84). Alexa-Fluor 555, 647 Phalloidins were used to label F-actin. CellLight-Talin was used to label focal adhesions in live cells. Blebbistatin was purchased from Millipore Sigma and used at a concentration of 5 μM. DAPI was used at 1:2000 to label cell nuclei.

### RNAi

AllStars Neg. Control siRNA (1027281) was from Qiagen, rat SUN1 (gene ID:360773, cat# M-080662-01-0005) and rat SUN2 (gene ID:315135, cat# M-081156-01-0005) smartpool siGENOME siRNAs, and MKL1 (gene ID:315151, cat# J-081405-08) ON-TARGETplus siRNA were from Dharmacon (Thermo Scientific). MTLn3 cells were transfected with siRNAs using Oligofectamine (Life Technologies) and optimum times for combined maximum KDs were found to be 24h (SUN1+SUN2+MKL1 siRNAs) or 48 h (SUN1+SUN2 siRNAs).

### PVA (polyvinyl alchohol) Film coating

The surface of glass substrates were coated with Polyvinyl Alcohol (PVA) in order to prevent cells from adhering to the 2D glass surface. This protocol was adapted from the literature for use with glass coverslips (64). Briefly, 12 mm glass coverslips were Piranha Cleaned (3:1, H2SO4:H2O2) for 10 minutes, rinsed 3X, and dried in N2. Coverslips were silanized in a container with 3-(amino)propyltrimethoxysilane (APTMS) at 65°C for 1 hr. Substrates were activated using glutaraldehyde for 30 min., rinsed 3X, and dried in N2. A 5.6% polyvinyl alcohol (PVA, Sigma-Aldrich, St. Louis, MO) solution was solubilized at 90°C, sonicated, and filtered through a 0.2 μm syringe filter. An 11.24% 2N HCl was added to the filtered solution. 65 μl of the PVA solution was added to each coverslip for 5 min. followed by spincoating at 7000 RPM for 40 sec. The coated coverslips were cured overnight at 4°C. A ring of polydimethylsiloxane (PDMS) was placed around the coverslip boundary in order to suspend the electrospun fibers over the PVA surface.

### Electrospun Fibers

One-dimensional aligned microfibers were created from a 21% (w/w) poly lactic-co-glycolic acid, PLGA (LACTEL) solution dissolved in hexafluoroisopropanol (HFIP, Krackeler) with 1% NaCl (Sigma Aldrich) and stirred overnight. The solution was fed into a 3 mL syringe, the flow rate was controlled by a syringe pump, and the high-voltage was applied using a high voltage power supply. The aligned microfibers were attracted to two grounded parallel collector plates consisting of cleaved silicon. A PVA coated glass coverslip was lifted between the parallel collector plates to transfer the electrospun fibers. The collector plates were 15cm from the needle tip. The flow rate was set to 16 μL/min and the voltage was set to 12 kV. The fibers were allowed to collect between the parallel plates for 6 seconds before the pump was stopped and the fibers were collected.

### ECM coating of fibers

The cover glass from a MatTek dish (10 mm well) was removed and the electrospun fiber containing 12 mm cover glass was attached to the MatTek dish using silicone glue. Alexa-568 labeld fibronectin (final conc = 10 ug/ml) or FITC labeled collagen I (final conc = 40 ug/ml) was added to the central well to completely cover the fibers. After 2 hours, fibers were washed 2X to remove the excess fluorescent ECM and dishes were stored in dark at 4 deg C. Labeled fibers were used within 1-2 days.

### Fibronectin-coated micro-patterned 1D lines

Custom made fibronectin-coated micro-patterned 1D lines and 2D areas (1D2D EDDY) were generated by CYTOO Inc. In addition, CYTOO chips Motility (CYTOO Inc.) was used as described (28). Dimensions of 1D lines were evaluated with AFM and showed a mean thickness of 2.5 μm and a depth of 3.5 nm (Fig S1).

### Scanning electron microscopy

Scanning electron microscopy was performed using a Leo1550 scanning electron microscope. This technique was used to measure microfibers to ensure that they were of the desired diameter. Fibers were coated by bombardment with gold-palladium particles in an argon gas environment. The chamber was pumped down to 50 mTorr and subsequently flooded with argon gas. The chamber was again pumped down to 50 mTorr and then coated with the gold-palladium particles. This process was used to reduce the charging effect of the specimen because of the high electrical conductivity of the gold-palladium coating. The in lens detector was used with the 20 μm aperture at 5 kV. The sizes were determined using Image J.

### Fibronectin staining of ECM in tumor sections

MTLn3 orthotopic tumors grown in SCID mice were removed and immediately frozen in liquid nitrogen. Tumor sections were cut for immunostaining. Tissue sections were blocked in donkey serum for 45 minutes and mouse anti-fibronectin antibody (Abcam, cat#6328) was added at 1:100 dilution for 1 hour. Sections were washed 3x in PBS and then incubated with a donkey anti-mouse Alexa 555 secondary antibody at 1:1000 dilution (Life technologies) for 1 hour. Secondary antibody was washed 3x with PBS. Images were obtained on a custom built multiphoton microscope described previously (85).

### Live-cell Microscopy

Time lapse images were acquired on a wide-field DeltaVision microscope equipped with a high precision X and Y NanoMotion III stage for capturing images at multiple positions and a Photometrics CoolSnap HQ2 CCD camera (Applied Precision, LLC). Imaging was done at 37°C with a 20× air objective, NA = 0.4. Up to 10-15 different fields along the linear dimension were imaged in the phase contrast and FITC channels every 2 min for up to 10-12 hours. For tumor cells and BACs co-migration experiment, MTLn3 cells were imaged alone for 2 h, followed by addition of BACs were added to the imaging chamber on the microscope stage and imaging was continued for the next 6-8 h. For tumor cell migration parameter calculations, we only analyzed tumor cells moving on a single fiber in the areas where fibers do not cross each other.

### Single Tumor Cell Tracking in 1D or 2D

A custom ImageJ macro was used to track single cells based on nucleus centroid tracking in GFP-H2B time-lapse movies (Fig. S3, movie 4). Only single non-dividing cells were tracked. Raw tracking values were used to calculate cell speed and persistence values as described earlier (28).

### Analysis of SUN1 and SUN2 expression in cancer patients

Expression of SUN1 and SUN2 in primary tumor vs normal breast tissue was performed on TCGA dataset for breast invasive carcinoma using UALCAN, http://ualcan.path.uab.edu/analysis.html (86). Analysis of SUN1 and SUN2 expression in tumor vs normal tissue across tumor types was performed using GEPIA, http://gepia.cancer-pku.cn (87).

### Statistical Analysis

All experiments were performed for at least 3 independent biological replicates. Statistical significance was calculated using unpaired, two-tailed Student’s t-test and statistical significance was defined as p value < 0.05. The * indicates p value <0.05; ** indicates p value <0.01; *** indicates p value <0.001 and error bars represent the SEM.

## SUPPLEMENTAL MOVIES

**Supplemental figure S1:**
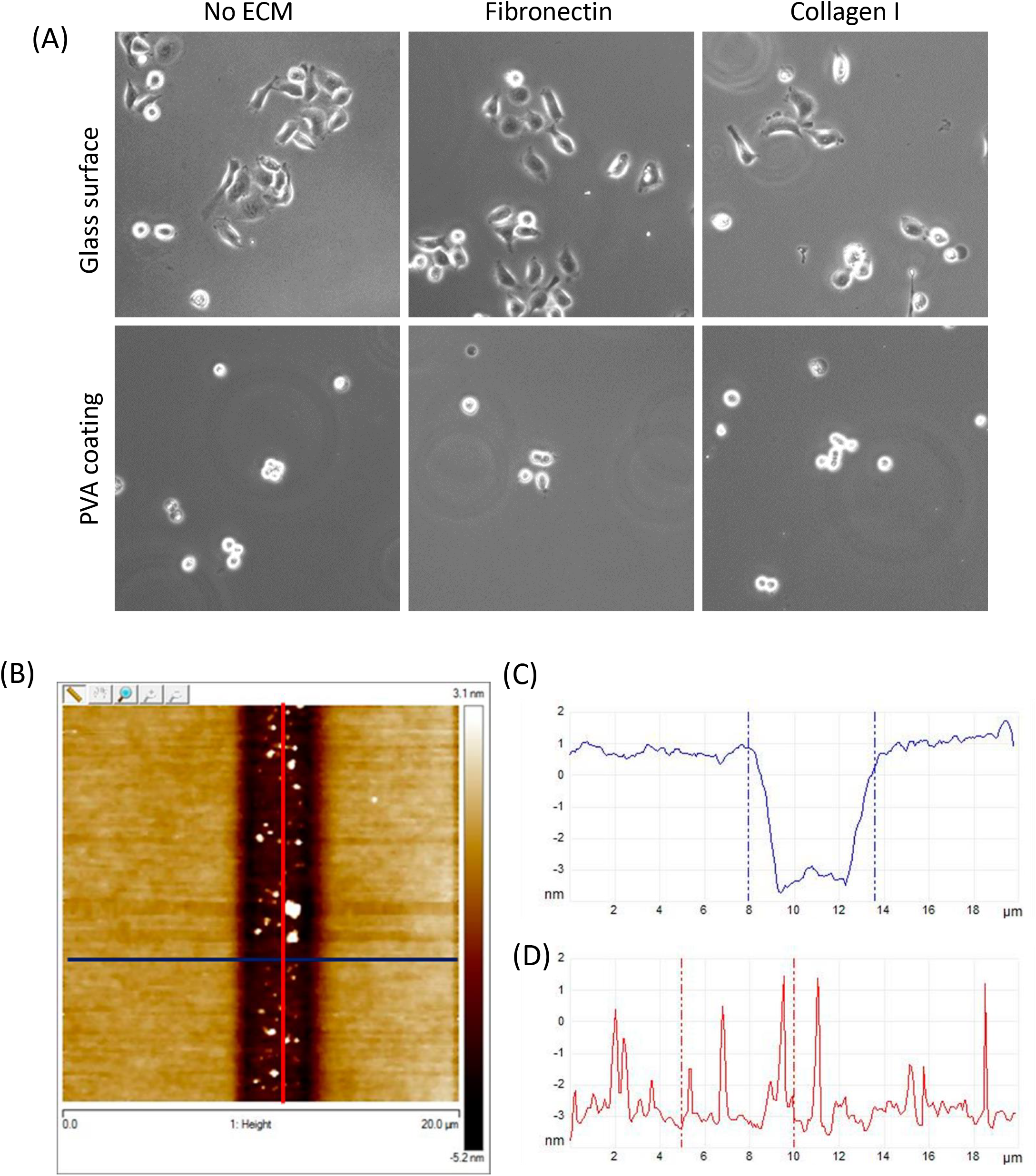
Surface characterization in 1D assays. (A) Tumor cells do not adhere and spread on PVA-coated surface. Images of MTLn3 cells plated on either glass or PVA-coated surface with and without ECM (fibronectin or collagen) coating. (B) The dimensions of 1D micro-patterned lines were evaluated using AFM. Image shows the height map of a 1D line and the surrounding area. (C) Height scan along the blue line in figure S1(B) shows that the thickness of 1D lines is approximately 2.5 μm. (D) Height scan along the red line in figure S1(B) shows that the depth of 1D lines is approximately 3-4 nm.

**Supplemental figure S2:**
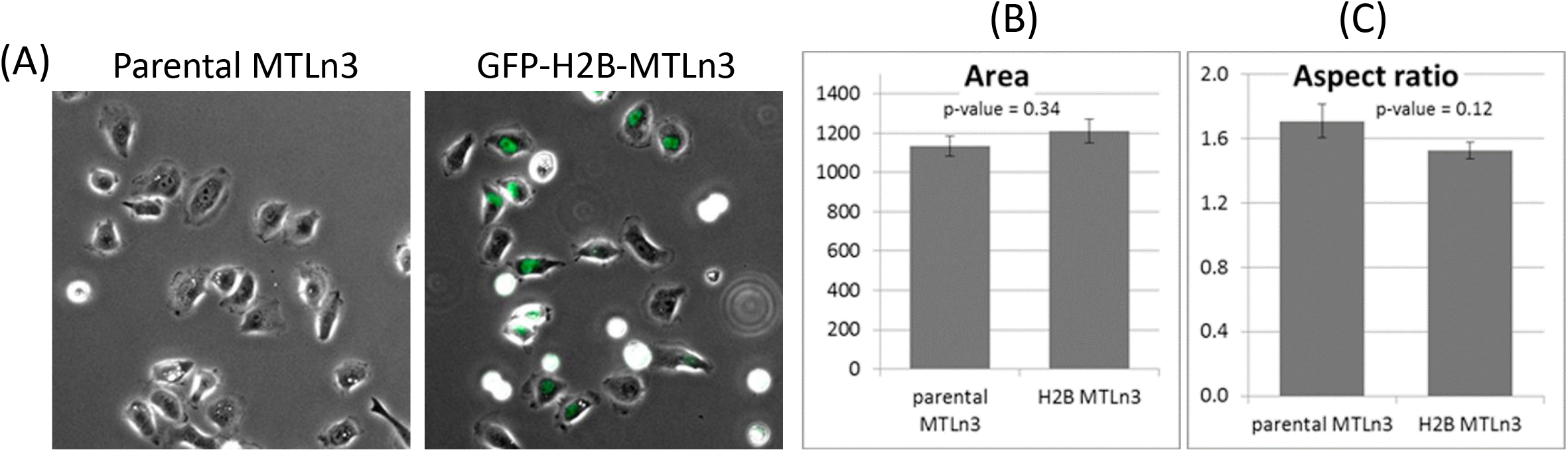
Cellular features of stable GFP-H2B-MTLn3 cells are similar to parental MTLn3 cells. (A) Parental or GFP-H2B expressing MTLn3 cells were plated on ECM coated 2D glass surfaces. Phase and phase+GFP channel images were acquired for the parental and GFP-H2B MTLn3 cells, respectively. (B, C) Quantifications of cell spread area (B) and cellular aspect ratio (C) in parental and GFP-H2B expressing MTLn3 cells.

**Supplemental figure S3:**
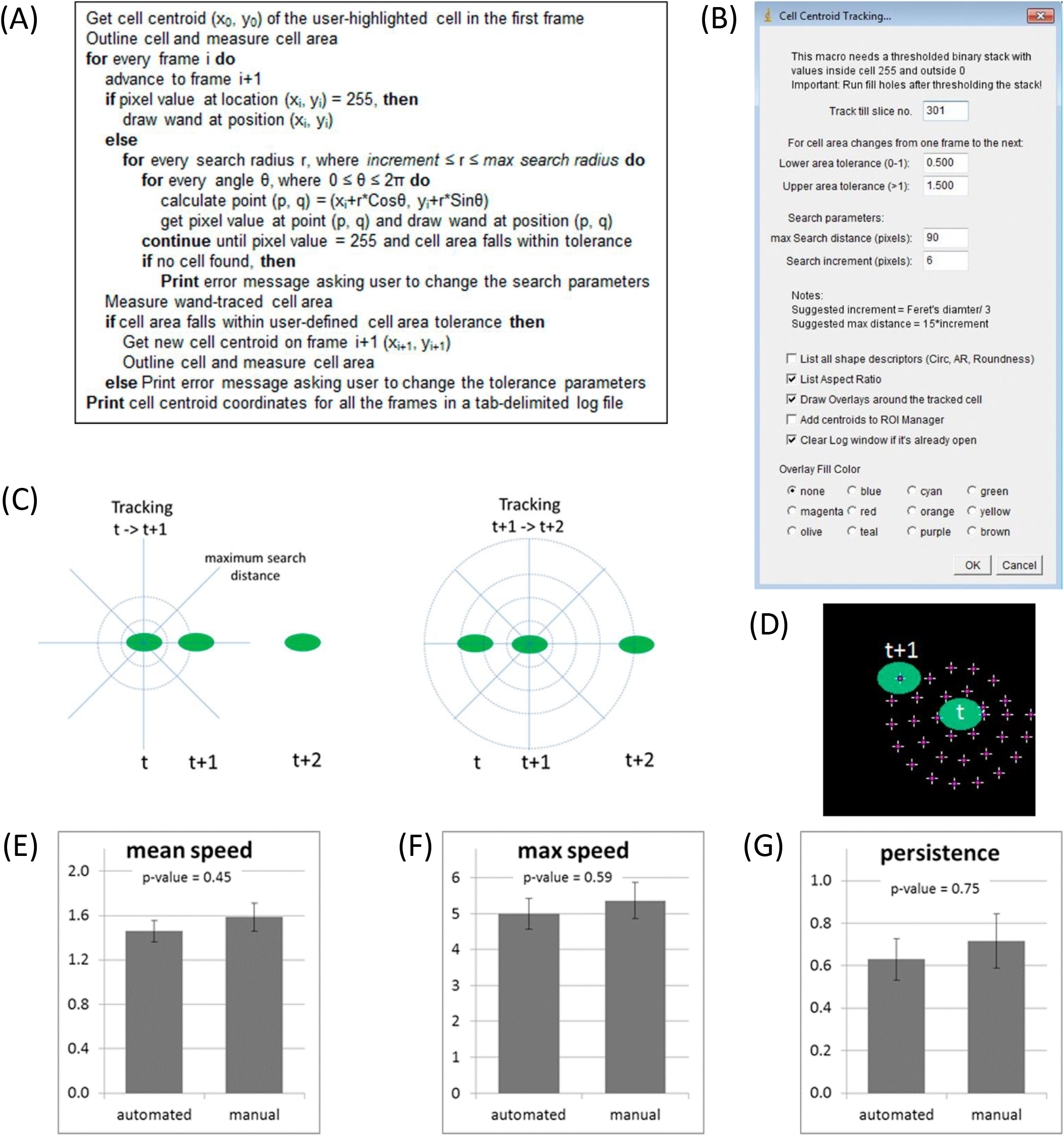
Description of “Cell centroid tracking” macro for automated tracking of single tumor cells. (A) Algorithm for cell centroid tracking macro. (B) Macro user-interface (UI) showing different parameters (e.g. maximum search distance, search increment etc.) which user can adjust for accurate cell tracking. (C) Schematic of cell nucleus tracking from time, t to t+1 and t+1 to t+2. Green ovals represent cell nucleus at different times and circles of different radii represent successive search increments. (D) An example of cell nucleus tracking from time, t to t+1. Green ovals represent cell nucleus at different time points and cross-hairs show all the points generated by the macro in search of the cell nucleus at time t+1. Search takes place in a clockwise manner. (E, F, G) GFP-H2B expressing tumor cells moving on 2 μm fibers were tracked using automated method (with Cell centroid tracking macro) and by marking cell centroid manually in each frame of the movie. Bar plots for the cell migration parameter calculations - average speed (E), maximum speed (F) and persistence (G), show that the automated method is as accurate as the manual method. Therefore, all subsequent single tumor cell speed and persistence calculations were performed using automated Cell centroid tracking macro.

**Supplemental figure S4:**
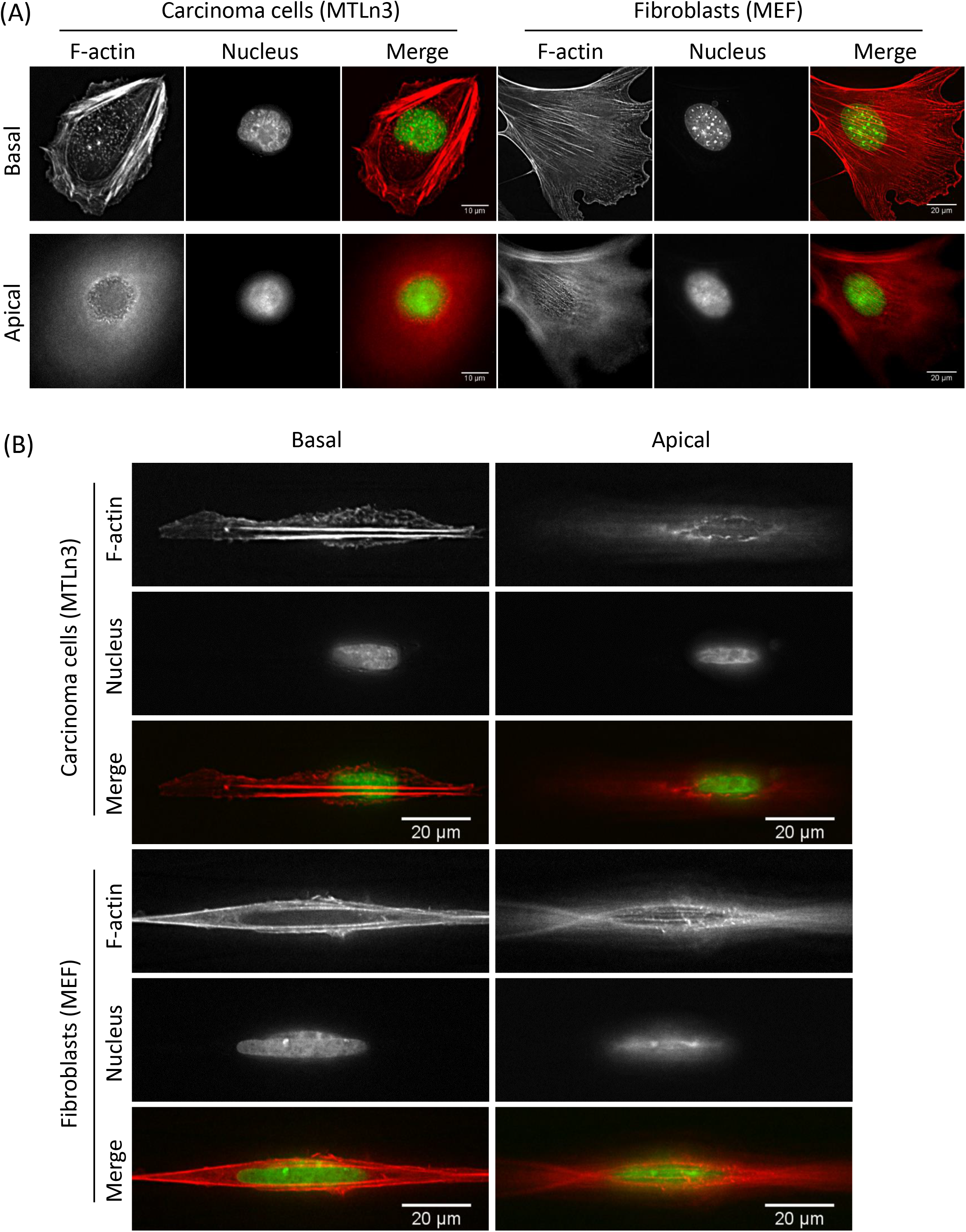
MTLn3 tumor cells do not display actin nuclear cap in 2D or 1D. (A) MTLn3 tumor cells and MEF fibroblasts were plated on 2D substrate and stained with phalloidin and DAPI. Top panels show basal plane and the bottom panels show apical plane from the z-stack. Note that MEFs have apical F-actin fibers which are missing in the MTLn3 cell. (B) MTLn3 tumor cells and MEF fibroblasts were plated on 1D substrate (2.5 μm micropatterned line) and stained with phalloidin and DAPI. Left panels show basal plane and the right panels show apical plane from the z-stack. Note that MEFs have apical F-actin fibers which are missing in the MTLn3 cell.

**Supplemental figure S5:**
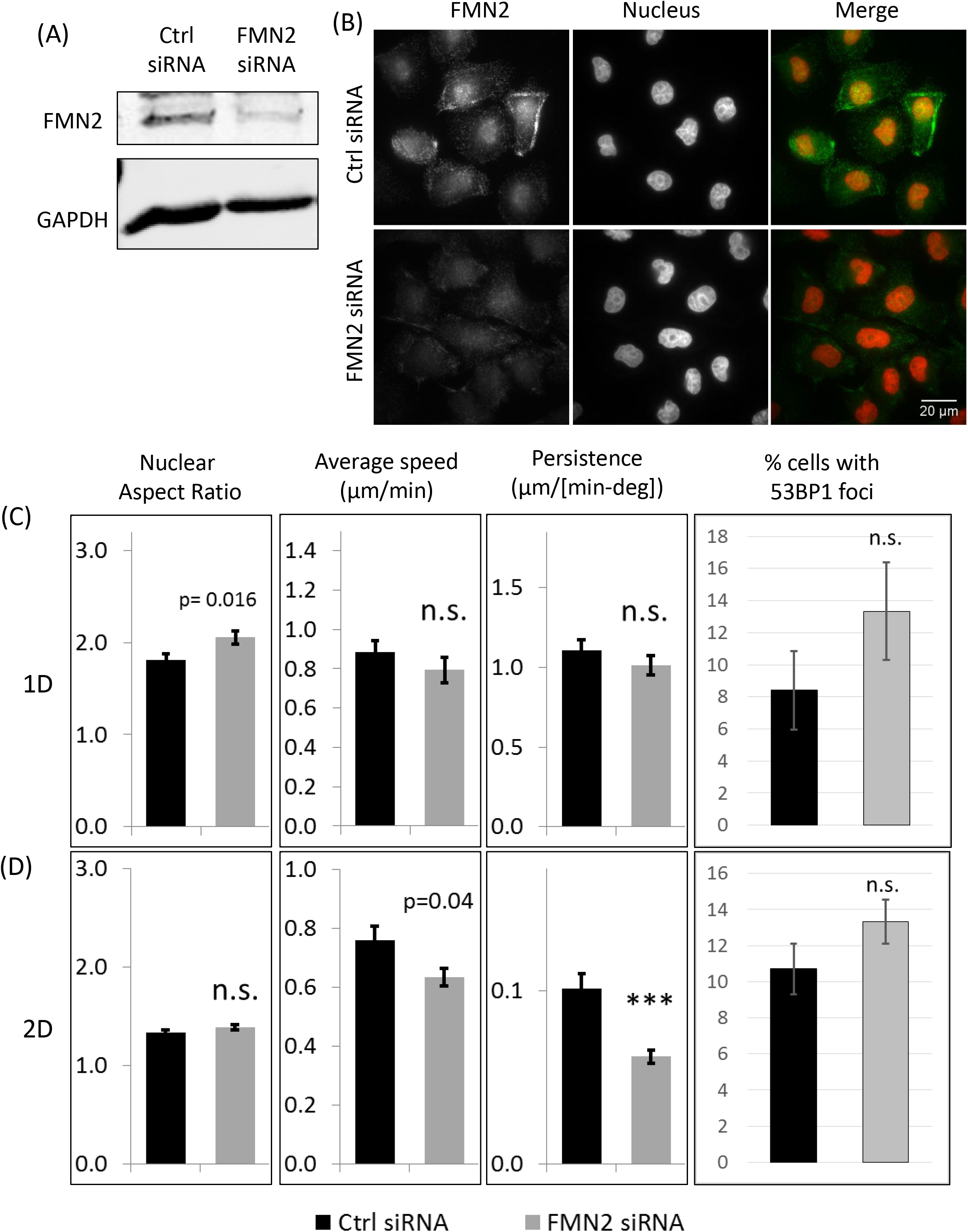
Formin FMN2 does not play any role in tumor cell motility in 1D. (A) Western blot showing FMN2 KD with FMN2 smartpool siRNA. (B) Immunofluorescence images of MTLn3 cells treated with control or FMN2 smartpool siRNA were stained with anti-FMN2 antibody. Please note the decrease in FMN2 fluorescence signal indicating efficient FMN2 KD. (C, D) Quantifications of nuclear aspect ratio, speed, persistence and % cells with 53BP1 foci in cells moving in 1D (C) or 2D (D).

**Supplemental figure S6:**
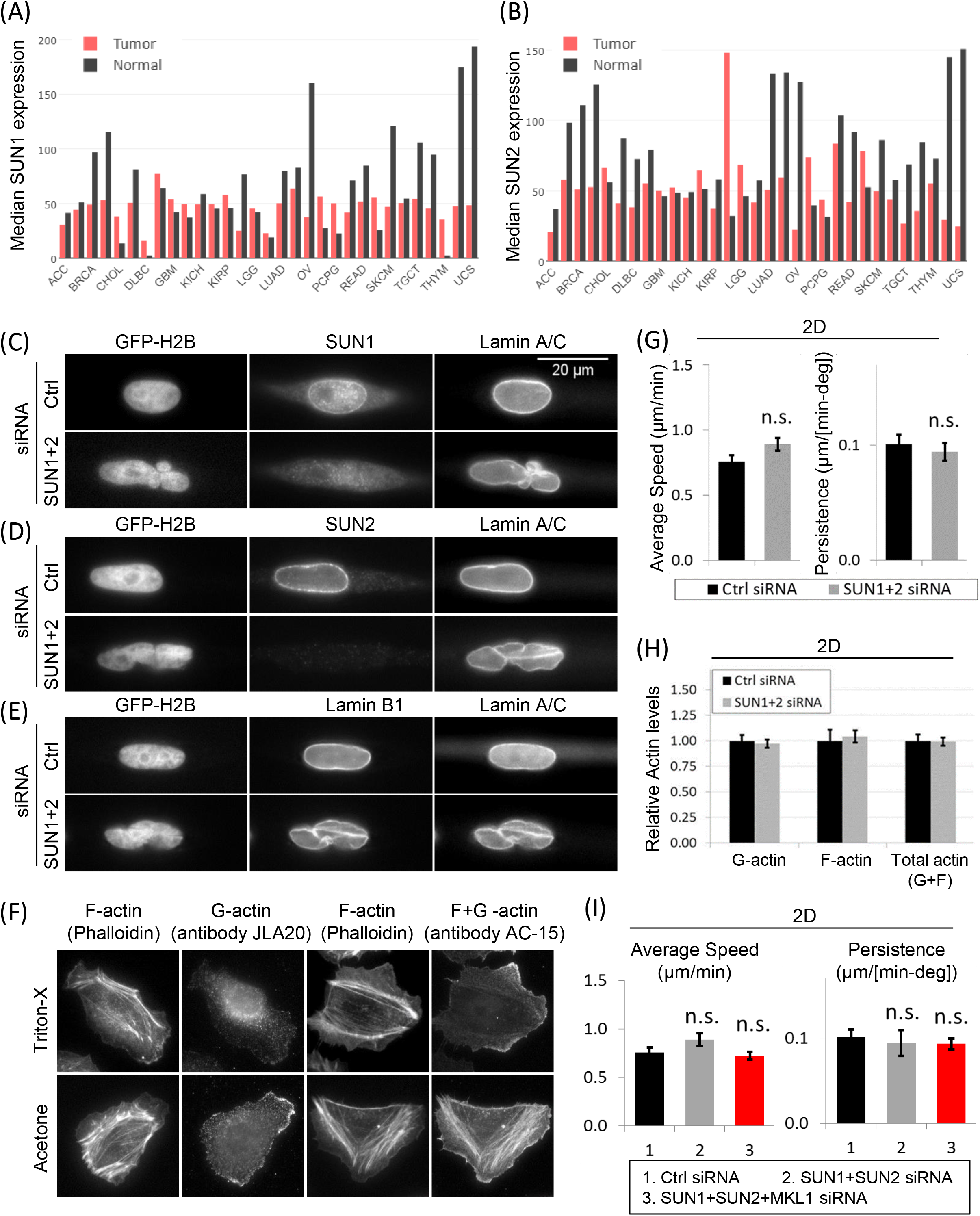
SUN1, SUN2 and MKL1 do not play any role in 2D tumor cell motility. (A, B) RNA-Seq data analysis for median SUN1 and SUN2 expression in tumor vs normal tissue across tumor types from the TCGA and GTEx datasets. Tumor type abbreviations - ACC: Adrenocortical carcinoma, BLCA: Bladder Urothelial Carcinoma, BRCA: Breast invasive carcinoma, CESC: Cervical squamous cell carcinoma and endocervical adenocarcinoma, CHOL: Cholangio carcinoma, COAD: Colon adenocarcinoma, DLBC: Lymphoid Neoplasm Diffuse Large B-cell Lymphoma, ESCA: Esophageal carcinoma, GBM: Glioblastoma multiforme, HNSC: Head and Neck squamous cell carcinoma, KICH: Kidney Chromophobe, KIRC: Kidney renal clear cell carcinoma, KIRP: Kidney renal papillary cell carcinoma, LAML: Acute Myeloid Leukemia, LGG: Brain Lower Grade Glioma, LIHC: Liver hepatocellular carcinoma, LUAD: Lung adenocarcinoma, LUSC: Lung squamous cell carcinoma, MESO: Mesothelioma, OV: Ovarian serous cystadenocarcinoma, PAAD: Pancreatic adenocarcinoma, PCPG: Pheochromocytoma and Paraganglioma, PRAD: Prostate adenocarcinoma, READ: Rectum adenocarcinoma, SARC: Sarcoma, SKCM: Skin Cutaneous Melanoma, STAD: Stomach adenocarcinoma, TGCT: Testicular Germ Cell Tumors, THCA: Thyroid carcinoma, THYM: Thymoma, UCEC: Uterine Corpus Endometrial Carcinoma, UCS: Uterine Carcinosarcoma, UVM: Uveal Melanoma. (C) Control and SUN1+2 siRNA treated tumor cells were stained with SUN1 and Lamin A/C antibodies. Immunofluorescence images show multi-lobular nucleus with no SUN1 localization at the nuclear envelope, whereas Lamin A/C localization was unchanged after SUN1+2 KD. (D) Control and SUN1+2 siRNA treated tumor cells were stained with SUN2 and Lamin A/C antibodies. Immunofluorescence images show multi-lobular nucleus with no SUN2 localization at the nuclear envelope, whereas Lamin A/C localization was unchanged after SUN1+2 KD. (E) Control and SUN1+2 siRNA treated tumor cells were stained with Lamin B1 and Lamin A/C antibodies. Immunofluorescence images show multi-lobular nucleus with no changes in Lamin B1 and Lamin A/C localizations after SUN1+2 KD. (F) Staining of tumor cells in 2D with phalloidin and either JLA antibody (specific for G-actin), or AC-15 antibody (labels both G-and F-actin) under Triton-X or acetone permeabilization conditions. Images show that acetone, but not Triton-X, permeabilization preserves the G-actin and F-actin architecture in the cells. (G) Quantifications of average tumor cell speed and persistence in control and SUN1+2 KD cells migrating in 2D. (H) Quantifications of whole cell F-actin, G-actin and total actin (F-actin + G-actin) levels in control and SUN1+2 KD cells migrating in 2D. (I) Quantifications of average tumor cell speed and tumor cell persistence after SUN1+SUN2+MKL1 KD in cells migrating in 2D. Control and SUN1+2 siRNA bars are shown here from figure S6G for comparison.

**Movie 1**: 3D reconstruction movies of collagen fibers *in vivo*, using second harmonic intravital imaging, in PyMT and MTLn3 tumor models

**Movie 2**: Movie showing tumor cells (GFP-H2B expressing MTLn3) moving on ECM coated 1D fibers. Upper panel shows phase contrast channel and lower panel shows GFP channel. Time in hour: minute.

**Movie 3**: Movie showing tumor cell (MTLn3) and macrophage (BAC) pairing and alternating streaming pattern on 1D fibers. Time in hour: minute.

**Movie 4:** Left: tumor cells moving in 2D. Right: tumor cells moving on 2.5 μm micro-patterned 1D lines. Upper panels show phase contrast channel. Lower panels show tracking of individual tumor cells based on GFP-H2B thresholding and automated tracking using Cell centroid tracking macro (Fig. S3). Different colors mark nuclei of different single tumor cells. Time in hour: minute.

**Movie 5:** GFP-talin expressing MTLn3 tumor cells moving on ECM coated 1D fiber. Time in hour: minute.

**Movie 6:** GFP-H2B expressing MTLn3 tumor cells moving on ECM coated 1D fibers, before and after 5 μM blebbistatin treatment. Time in hour: minute.

**Movie 7**: GFP-H2B expressing MTLn3 tumor cells moving on ECM coated 1D fiber. Time in hour: minute.

**Movie 8**: GFP-H2B expressing MTLn3 tumor cells moving on ECM coated 2D surface. Time in hour: minute.

**Movie 9:** Intravital imaging of GFP-H2B expressing MTLn3 tumor cells moving *in vivo*.

**Movie 10:** GFP-H2B expressing MTLn3 tumor cells treated with SUN1+2 siRNA were plated on 2D substrate and time-lapse images for phase and GFP-H2B channels were recorded every 2 min. Time in hour: minute.

**Movie 11:** GFP-H2B expressing MTLn3 tumor cells treated with SUN1+2 siRNA were plated on 2.5 μm, 1D micro-patterned lines and time-lapse images for phase and GFP-H2B channels were recorded every 2 min. Time in hour: minute.

## REFERENCES

1. Joyce JA, Pollard JW. (2009). Microenvironmental regulation of metastasis. Nature reviews Cancer. 9(4):239–52. PMID: 19279573 / PMCID: PMC3251309.

2. Place AE, Jin Huh S, Polyak K. (2011). The microenvironment in breast cancer progression: biology and implications for treatment. Breast cancer research : BCR. 13(6):227. PMID: 22078026 / PMCID: PMCPMC3326543.

3. Robinson BD, Sica GL, Liu YF, Rohan TE, Gertler FB, Condeelis JS, Jones JG. (2009). Tumor microenvironment of metastasis in human breast carcinoma: a potential prognostic marker linked to hematogenous dissemination. Clin Cancer Res. 15(7):2433–41. PMID: 19318480 / PMCID: PMCPMC3156570.

4. Roussos ET, Condeelis JS, Patsialou A. (2011). Chemotaxis in cancer. Nature reviews Cancer. 11(8):573–87. PMID: 21779009 / PMCID: PMC4030706.

5. Sidani M, Wyckoff J, Xue C, Segall JE, Condeelis J. (2006). Probing the microenvironment of mammary tumors using multiphoton microscopy. J Mammary Gland Biol Neoplasia. 11(2):151–63. PMID: 17106644.

6. Balkwill FR, Capasso M, Hagemann T. (2012). The tumor microenvironment at a glance. Journal of Cell Science. 125(Pt 23):5591-6. PMID: 23420197.

7. Condeelis J, Pollard JW. (2006). Macrophages: obligate partners for tumor cell migration, invasion, and metastasis. Cell. 124(2):263–6. PMID: 16439202.

8. Quail DF, Joyce JA. (2013). Microenvironmental regulation of tumor progression and metastasis. Nat Med. 19(11):1423–37. PMID: 24202395 / PMCID: PMC3954707.

9. Alowami S, Troup S, Al-Haddad S, Kirkpatrick I, Watson PH. (2003). Mammographic density is related to stroma and stromal proteoglycan expression. Breast cancer research : BCR. 5(5):R129–35. PMID: 12927043 / PMCID: PMCPMC314426.

10. Guo YP, Martin LJ, Hanna W, Banerjee D, Miller N, Fishell E, Khokha R, Boyd NF. (2001). Growth factors and stromal matrix proteins associated with mammographic densities. Cancer Epidemiol Biomarkers Prev. 10(3):243–8. PMID: 11303594.

11. Provenzano PP, Eliceiri KW, Campbell JM, Inman DR, White JG, Keely PJ. (2006). Collagen reorganization at the tumor-stromal interface facilitates local invasion. BMC medicine. 4(1):38. PMID: 17190588 / PMCID: PMC1781458.

12. Moreau V, Saltel F. (2015). Type I collagen fibrils and discoidin domain receptor 1 set invadosomes straight. Mol Cell Oncol. 2(4):e1004963. PMID: 27308500 / PMCID: PMC4905343.

13. Provenzano PP, Inman DR, Eliceiri KW, Knittel JG, Yan L, Rueden CT, White JG, Keely PJ. (2008). Collagen density promotes mammary tumor initiation and progression. BMC medicine. 6:11. PMID: 18442412 / PMCID: PMC2386807.

14. Ramaswamy S, Ross KN, Lander ES, Golub TR. (2003). A molecular signature of metastasis in primary solid tumors. Nat Genet. 33(1):49–54. PMID: 12469122.

15. Conklin MW, Eickhoff JC, Riching KM, Pehlke CA, Eliceiri KW, Provenzano PP, Friedl A, Keely PJ. (2011). Aligned collagen is a prognostic signature for survival in human breast carcinoma. The American journal of pathology. 178(3):1221–32. PMID: 21356373 / PMCID: PMC3070581.

16. Han W, Chen S, Yuan W, Fan Q, Tian J, Wang X, Chen L, Zhang X, Wei W, Liu R, Qu J, Jiao Y, Austin RH, Liu L. (2016). Oriented collagen fibers direct tumor cell intravasation. Proc Natl Acad Sci U S A. 113(40):11208–13. PMID: 27663743 / PMCID: PMCPMC5056065.

17. Nelson MT, Short A, Cole SL, Gross AC, Winter J, Eubank TD, Lannutti JJ. (2014). Preferential, enhanced breast cancer cell migration on biomimetic electrospun nanofiber ‘cell highways’. BMC Cancer. 14:825. PMID: 25385001 / PMCID: PMCPMC4236463.

18. Oudin MJ, Jonas O, Kosciuk T, Broye LC, Guido BC, Wyckoff J, Riquelme D, Lamar JM, Asokan SB, Whittaker C, Ma D, Langer R, Cima MJ, Wisinski KB, Hynes RO, Lauffenburger DA, Keely PJ, Bear JE, Gertler FB. (2016). Tumor Cell-Driven Extracellular Matrix Remodeling Drives Haptotaxis during Metastatic Progression. Cancer Discov. 6(5):516–31. PMID: 26811325 / PMCID: PMC4854754.

19. Egeblad M, Rasch MG, Weaver VM. (2010). Dynamic interplay between the collagen scaffold and tumor evolution. Curr Opin Cell Biol. 22(5):697–706. PMID: 20822891 / PMCID: PMCPMC2948601.

20. Levental KR, Yu H, Kass L, Lakins JN, Egeblad M, Erler JT, Fong SF, Csiszar K, Giaccia A, Weninger W, Yamauchi M, Gasser DL, Weaver VM. (2009). Matrix crosslinking forces tumor progression by enhancing integrin signaling. Cell. 139(5):891–906. PMID: 19931152 / PMCID: PMC2788004.

21. Park CC, Rembert J, Chew K, Moore D, Kerlikowske K. (2009). High mammographic breast density is independent predictor of local but not distant recurrence after lumpectomy and radiotherapy for invasive breast cancer. Int J Radiat Oncol Biol Phys. 73(1):75–9. PMID: 18692323.

22. Riching KM, Cox BL, Salick MR, Pehlke C, Riching AS, Ponik SM, Bass BR, Crone WC, Jiang Y, Weaver AM, Eliceiri KW, Keely PJ. (2014). 3D collagen alignment limits protrusions to enhance breast cancer cell persistence. Biophys J. 107(11):2546–58. PMID: 25468334 / PMCID: PMCPMC4255204.

23. Gligorijevic B, Bergman A, Condeelis J. (2014). Multiparametric classification links tumor microenvironments with tumor cell phenotype. PLoS Biol. 12(11):e1001995. PMID: 25386698 / PMCID: PMC4227649

24. Condeelis J, Segall JE. (2003). Intravital imaging of cell movement in tumours. Nature reviews Cancer. 3(12):921–30. PMID: 14737122.

25. Wang W, Wyckoff JB, Frohlich VC, Oleynikov Y, Huttelmaier S, Zavadil J, Cermak L, Bottinger EP, Singer RH, White JG, Segall JE, Condeelis JS. (2002). Single cell behavior in metastatic primary mammary tumors correlated with gene expression patterns revealed by molecular profiling. Cancer Res. 62(21):6278–88. PMID: 12414658.

26. Wyckoff JB, Wang Y, Lin EY, Li JF, Goswami S, Stanley ER, Segall JE, Pollard JW, Condeelis J. (2007). Direct visualization of macrophage-assisted tumor cell intravasation in mammary tumors. Cancer Res. 67(6):2649–56. PMID: 17363585.

27. Xue C, Wyckoff J, Liang F, Sidani M, Violini S, Tsai KL, Zhang ZY, Sahai E, Condeelis J, Segall JE. (2006). Epidermal growth factor receptor overexpression results in increased tumor cell motility in vivo coordinately with enhanced intravasation and metastasis. Cancer Res. 66(1):192–7. PMID: 16397232.

28. Sharma VP, Beaty BT, Patsialou A, Liu H, Clarke M, Cox D, Condeelis JS, Eddy RJ. (2012). Reconstitution of in vivo macrophage-tumor cell pairing and streaming motility on one-dimensional micro-patterned substrates. Intravital. 1(1):77–85. PMID: 24634804 / PMCID: PMC3908597.

29. Sharma VP, Beaty BT, Cox D, Condeelis JS, Eddy RJ. (2014). An in vitro one-dimensional assay to study growth factor-regulated tumor cell-macrophage interaction. Methods in molecular biology. 1172:115-23. PMID: 24908299 / PMCID: PMC4124813.

30. Patsialou A, Bravo-Cordero JJ, Wang Y, Entenberg D, Liu H, Clarke M, Condeelis JS. (2013). Intravital multiphoton imaging reveals multicellular streaming as a crucial component of in vivo cell migration in human breast tumors. Intravital. 2(2):e25294. PMID: 25013744 / PMCID: PMC3908591.

31. Tong Z, Balzer EM, Dallas MR, Hung WC, Stebe KJ, Konstantopoulos K. (2012). Chemotaxis of cell populations through confined spaces at single-cell resolution. PLoS One. 7(1):e29211. PMID: 22279529 / PMCID: PMCPMC3261140.

32. Petrie RJ, Yamada KM. (2012). At the leading edge of three-dimensional cell migration. Journal of Cell Science. 125(Pt 24):5917-26. PMID: 23378019 / PMCID: PMCPMC4067260.

33. Friedl P, Wolf K. (2010). Plasticity of cell migration: a multiscale tuning model. J Cell Biol. 188(1):11–9. PMID: 19951899 / PMCID: PMCPMC2812848.

34. Doyle AD, Wang FW, Matsumoto K, Yamada KM. (2009). One-dimensional topography underlies three-dimensional fibrillar cell migration. J Cell Biol. 184(4):481–90. PMID: 19221195/ PMCID: PMCPMC2654121.

35. Lin B, Yin T, Wu YI, Inoue T, Levchenko A. (2015). Interplay between chemotaxis and contact inhibition of locomotion determines exploratory cell migration. Nat Commun. 6:6619. PMID: 25851023 / PMCID: PMCPMC4391292.

36. Martin K, Vilela M, Jeon NL, Danuser G, Pertz O. (2014). A growth factor-induced, spatially organizing cytoskeletal module enables rapid and persistent fibroblast migration. Dev Cell. 30(6):701–16. PMID: 25268172 / PMCID: PMCPMC4385272.

37. Paul CD, Hung WC, Wirtz D, Konstantopoulos K. (2016). Engineered Models of Confined Cell Migration. Annu Rev Biomed Eng. 18:159-80. PMID: 27420571.

38. Sheets K, Wunsch S, Ng C, Nain AS. (2013). Shape-dependent cell migration and focal adhesion organization on suspended and aligned nanofiber scaffolds. Acta Biomater. 9(7):7169–77. PMID: 23567946.

39. Fraley SI, Wu PH, He L, Feng Y, Krisnamurthy R, Longmore GD, Wirtz D. (2015). Three-dimensional matrix fiber alignment modulates cell migration and MT1-MMP utility by spatially and temporally directing protrusions. Sci Rep. 5:14580. PMID: 26423227 / PMCID: PMCPMC4589685.

40. Friedl P, Wolf K, Lammerding J. (2011). Nuclear mechanics during cell migration. Curr Opin Cell Biol. 23(1):55–64. PMID: 21109415 / PMCID: PMC3073574.

41. Khatau SB, Hale CM, Stewart-Hutchinson PJ, Patel MS, Stewart CL, Searson PC, Hodzic D, Wirtz D. (2009). A perinuclear actin cap regulates nuclear shape. Proc Natl Acad Sci U S A. 106(45):19017–22. PMID: 19850871 / PMCID: PMCPMC2776434.

42. Kim DH, Cho S, Wirtz D. (2014). Tight coupling between nucleus and cell migration through the perinuclear actin cap. Journal of Cell Science. 127(Pt 11):2528-41. PMID: 24639463 / PMCID: PMCPMC4038945.

43. Skau CT, Fischer RS, Gurel P, Thiam HR, Tubbs A, Baird MA, Davidson MW, Piel M, Alushin GM, Nussenzweig A, Steeg PS, Waterman CM. (2016). FMN2 Makes Perinuclear Actin to Protect Nuclei during Confined Migration and Promote Metastasis. Cell. 167(6):1571–85.e18. PMID: 27839864 / PMCID: PMCPMC5135586.

44. Denais C, Lammerding J. (2014). Nuclear mechanics in cancer. Advances in experimental medicine and biology. 773:435-70. PMID: 24563360 / PMCID: PMCPMC4591936.

45. Lombardi ML, Jaalouk DE, Shanahan CM, Burke B, Roux KJ, Lammerding J. (2011). The interaction between nesprins and sun proteins at the nuclear envelope is critical for force transmission between the nucleus and cytoskeleton. The Journal of biological chemistry. 286(30):26743–53. PMID: 21652697 / PMCID: PMC3143636.

46. King SJ, Nowak K, Suryavanshi N, Holt I, Shanahan CM, Ridley AJ. (2014). Nesprin-1 and nesprin-2 regulate endothelial cell shape and migration. Cytoskeleton (Hoboken). 71(7):423–34. PMID: 24931616.

47. Chancellor TJ, Lee J, Thodeti CK, Lele T. (2010). Actomyosin tension exerted on the nucleus through nesprin-1 connections influences endothelial cell adhesion, migration, and cyclic strain-induced reorientation. Biophys J. 99(1):115–23. PMID: 20655839 / PMCID: PMCPMC2895377.

48. Padmakumar VC, Libotte T, Lu W, Zaim H, Abraham S, Noegel AA, Gotzmann J, Foisner R, Karakesisoglou I. (2005). The inner nuclear membrane protein Sun1 mediates the anchorage of Nesprin-2 to the nuclear envelope. Journal of Cell Science. 118(Pt 15):3419-30. PMID: 16079285.

49. Crisp M, Liu Q, Roux K, Rattner JB, Shanahan C, Burke B, Stahl PD, Hodzic D. (2006). Coupling of the nucleus and cytoplasm: role of the LINC complex. J Cell Biol. 172(1):41–53. PMID: 16380439 / PMCID: PMCPMC2063530.

50. Kirby TJ, Lammerding J. (2018). Emerging views of the nucleus as a cellular mechanosensor. Nature cell biology. 20(4):373–81. PMID: 29467443 / PMCID: PMCPMC6440800.

51. Dahl KN, Ribeiro AJ, Lammerding J. (2008). Nuclear shape, mechanics, and mechanotransduction. Circulation research. 102(11):1307–18. PMID: 18535268 / PMCID: PMC2717705.

52. Matsumoto A, Hieda M, Yokoyama Y, Nishioka Y, Yoshidome K, Tsujimoto M, Matsuura N. (2015). Global loss of a nuclear lamina component, lamin A/C, and LINC complex components SUN1, SUN2, and nesprin-2 in breast cancer. Cancer Med. 4(10):1547–57. PMID: 26175118 / PMCID: PMC4618625.

53. Olson EN, Nordheim A. (2010). Linking actin dynamics and gene transcription to drive cellular motile functions. Nat Rev Mol Cell Biol. 11(5):353–65. PMID: 20414257 / PMCID: PMCPMC3073350.

54. Finch-Edmondson M, Sudol M. (2016). Framework to function: mechanosensitive regulators of gene transcription. Cell Mol Biol Lett. 21:28. PMID: 28536630 / PMCID: PMCPMC5415767.

55. Vartiainen MK, Guettler S, Larijani B, Treisman R. (2007). Nuclear actin regulates dynamic subcellular localization and activity of the SRF cofactor MAL. Science. 316(5832):1749–52. PMID: 17588931.

56. Lu P, Takai K, Weaver VM, Werb Z. (2011). Extracellular matrix degradation and remodeling in development and disease. Cold Spring Harbor perspectives in biology. 3(12). PMID: 21917992 / PMCID: PMCPMC3225943.

57. Hall G, Liang W, Li X. (2017). Fitting-free algorithm for efficient quantification of collagen fiber alignment in SHG imaging applications. Biomed Opt Express. 8(10):4609–20. PMID: 29082088 / PMCID: PMCPMC5654803.

58. Kakkad SM, Solaiyappan M, Argani P, Sukumar S, Jacobs LK, Leibfritz D, Bhujwalla ZM, Glunde K. (2012). Collagen I fiber density increases in lymph node positive breast cancers: pilot study. Journal of biomedical optics. 17(11):116017. PMID: 23117811 / PMCID: PMCPMC3486274.

59. Kakkad SM, Solaiyappan M, O’Rourke B, Stasinopoulos I, Ackerstaff E, Raman V, Bhujwalla ZM, Glunde K. (2010). Hypoxic tumor microenvironments reduce collagen I fiber density. Neoplasia. 12(8):608–17. PMID: 20689755 / PMCID: PMCPMC2915405.

60. Hulmes DJ. (2002). Building collagen molecules, fibrils, and suprafibrillar structures. J Struct Biol. 137(1–2):2-10. PMID: 12064927.

61. Parry DA, Barnes GR, Craig AS. (1978). A comparison of the size distribution of collagen fibrils in connective tissues as a function of age and a possible relation between fibril size distribution and mechanical properties. Proc R Soc Lond B Biol Sci. 203(1152):305–21. PMID: 33395.

62. Yevick HG, Duclos G, Bonnet I, Silberzan P. (2015). Architecture and migration of an epithelium on a cylindrical wire. Proc Natl Acad Sci U S A. 112(19):5944–9. PMID: 25922533 / PMCID: PMCPMC4434757.

63. Meehan S, Nain AS. (2014). Role of suspended fiber structural stiffness and curvature on single-cell migration, nucleus shape, and focal-adhesion-cluster length. Biophys J. 107(11):2604–11. PMID: 25468339 / PMCID: PMCPMC4255195.

64. Doyle AD, Wang FW, Matsumoto K, Yamada KM. (2009). One-dimensional topography underlies three-dimensional fibrillar cell migration. Journal of Cell Biology. 184(4):481–90. PMID: 19221195 / PMCID: PMC2654121.

65. Qin S, Ricotta V, Simon M, Clark RA, Rafailovich MH. (2015). Continual cell deformation induced via attachment to oriented fibers enhances fibroblast cell migration. PLoS One. 10(3):e0119094. PMID: 25774792 / PMCID: PMCPMC4361054.

66. Versaevel M, Grevesse T, Gabriele S. (2012). Spatial coordination between cell and nuclear shape within micropatterned endothelial cells. Nat Commun. 3:671. PMID: 22334074.

67. Wolf K, Te Lindert M, Krause M, Alexander S, Te Riet J, Willis AL, Hoffman RM, Figdor CG, Weiss SJ, Friedl P. (2013). Physical limits of cell migration: control by ECM space and nuclear deformation and tuning by proteolysis and traction force. J Cell Biol. 201(7):1069–84. PMID: 23798731 / PMCID: PMC3691458.

68. Thomas DG, Yenepalli A, Denais CM, Rape A, Beach JR, Wang YL, Schiemann WP, Baskaran H, Lammerding J, Egelhoff TT. (2015). Non-muscle myosin IIB is critical for nuclear translocation during 3D invasion. J Cell Biol. 210(4):583–94. PMID: 26261182 / PMCID: PMC4539979.

69. Davidson PM, Denais C, Bakshi MC, Lammerding J. (2014). Nuclear deformability constitutes a rate-limiting step during cell migration in 3-D environments. Cell Mol Bioeng. 7(3):293–306. PMID: 25436017 / PMCID: PMCPMC4243304.

70. Krause M, Yang FW, Te Lindert M, Isermann P, Schepens J, Maas RJA, Venkataraman C, Lammerding J, Madzvamuse A, Hendriks W, Te Riet J, Wolf K. (2019). Cell migration through three-dimensional confining pores: speed accelerations by deformation and recoil of the nucleus. Philos Trans R Soc Lond B Biol Sci. 374(1779):20180225. PMID: 31431171 / PMCID: PMCPMC6627020.

71. Yamauchi K, Yang M, Jiang P, Yamamoto N, Xu M, Amoh Y, Tsuji K, Bouvet M, Tsuchiya H, Tomita K, Moossa AR, Hoffman RM. (2005). Real-time in vivo dual-color imaging of intracapillary cancer cell and nucleus deformation and migration. Cancer Res. 65(10):4246–52. PMID: 15899816.

72. Jain N, Iyer KV, Kumar A, Shivashankar GV. (2013). Cell geometric constraints induce modular gene-expression patterns via redistribution of HDAC3 regulated by actomyosin contractility. Proc Natl Acad Sci U S A. 110(28):11349–54. PMID: 23798429 / PMCID: PMCPMC3710882.

73. Harada T, Swift J, Irianto J, Shin JW, Spinler KR, Athirasala A, Diegmiller R, Dingal PC, Ivanovska IL, Discher DE. (2014). Nuclear lamin stiffness is a barrier to 3 D migration, but softness can limit survival. J Cell Biol. 204(5):669–82. PMID: 24567359 / PMCID: PMC3941057.

74. Swaminathan V, Mythreye K, O’Brien ET, Berchuck A, Blobe GC, Superfine R. (2011). Mechanical stiffness grades metastatic potential in patient tumor cells and in cancer cell lines. Cancer Res. 71(15):5075–80. PMID: 21642375 / PMCID: PMCPMC3220953.

75. Suresh S. (2007). Nanomedicine: elastic clues in cancer detection. Nat Nanotechnol. 2(12):748–9. PMID: 18654425.

76. Guck J, Schinkinger S, Lincoln B, Wottawah F, Ebert S, Romeyke M, Lenz D, Erickson HM, Ananthakrishnan R, Mitchell D, Kas J, Ulvick S, Bilby C. (2005). Optical deformability as an inherent cell marker for testing malignant transformation and metastatic competence. Biophys J. 88(5):3689–98. PMID: 15722433 / PMCID: PMCPMC1305515.

77. Cross SE, Jin YS, Rao J, Gimzewski JK. (2007). Nanomechanical analysis of cells from cancer patients. Nat Nanotechnol. 2(12):780–3. PMID: 18654431.

78. Lee MH, Wu PH, Staunton JR, Ros R, Longmore GD, Wirtz D. (2012). Mismatch in mechanical and adhesive properties induces pulsating cancer cell migration in epithelial monolayer. Biophys J. 102(12):2731–41. PMID: 22735523 / PMCID: PMCPMC3379010.

79. Zwerger M, Ho CY, Lammerding J. (2011). Nuclear mechanics in disease. Annu Rev Biomed Eng. 13:397-428. PMID: 21756143 / PMCID: PMCPMC4600467.

80. Zink D, Fischer AH, Nickerson JA. (2004). Nuclear structure in cancer cells. Nature reviews Cancer. 4(9):677–87. PMID: 15343274.

81. Lammerding J, Fong LG, Ji JY, Reue K, Stewart CL, Young SG, Lee RT. (2006). Lamins A and C but not lamin B1 regulate nuclear mechanics. The Journal of biological chemistry. 281(35):25768–80. PMID: 16825190.

82. Neelam S, Chancellor TJ, Li Y, Nickerson JA, Roux KJ, Dickinson RB, Lele TP. (2015). Direct force probe reveals the mechanics of nuclear homeostasis in the mammalian cell. Proc Natl Acad Sci U S A. 112(18):5720–5. PMID: 25901323 / PMCID: PMCPMC4426403.

83. Gau D, Roy P. (2018). SRF’ing and SAP’ing - the role of MRTF proteins in cell migration. Journal of Cell Science. 131(19). PMID: 30309957 / PMCID: PMCPMC6919568.

84. Lee CW, Vitriol EA, Shim S, Wise AL, Velayutham RP, Zheng JQ. (2013). Dynamic localization of G-actin during membrane protrusion in neuronal motility. Curr Biol. 23(12):1046–56. PMID: 23746641 / PMCID: PMCPMC3712510.

85. Entenberg D, Wyckoff J, Gligorijevic B, Roussos ET, Verkhusha VV, Pollard JW, Condeelis J. (2011). Setup and use of a two-laser multiphoton microscope for multichannel intravital fluorescence imaging. Nat Protoc. 6(10):1500–20. PMID: 21959234 / PMCID: PMC4028841.

86. Chandrashekar DS, Bashel B, Balasubramanya SAH, Creighton CJ, Ponce-Rodriguez I, Chakravarthi B, Varambally S. (2017). UALCAN: A Portal for Facilitating Tumor Subgroup Gene Expression and Survival Analyses. Neoplasia. 19(8):649–58. PMID: 28732212 / PMCID: PMCPMC5516091.

87. Tang Z, Li C, Kang B, Gao G, Li C, Zhang Z. (2017). GEPIA: a web server for cancer and normal gene expression profiling and interactive analyses. Nucleic Acids Res. 45(W1):W98–w102. PMID: 28407145 / PMCID: PMCPMC5570223.

88. Burridge K, Wittchen ES. (2013). The tension mounts: stress fibers as force-generating mechanotransducers. J Cell Biol. 200(1):9–19. PMID: 23295347 / PMCID: PMCPMC3542796.

89. Balaban NQ, Schwarz US, Riveline D, Goichberg P, Tzur G, Sabanay I, Mahalu D, Safran S, Bershadsky A, Addadi L, Geiger B. (2001). Force and focal adhesion assembly: a close relationship studied using elastic micropatterned substrates. Nature cell biology. 3(5):466–72. PMID: 11331874.

